# Evolution of dynamical networks enhances catalysis in a designer enzyme

**DOI:** 10.1101/2020.08.21.260885

**Authors:** H. Adrian Bunzel, J. L. Ross Anderson, Donald Hilvert, Vickery L. Arcus, Marc W. van der Kamp, Adrian J. Mulholland

## Abstract

Activation heat capacity is emerging as a crucial factor in enzyme thermoadaptation, as shown by non-Arrhenius behaviour of many natural enzymes^1,2^. However, its physical origin and relationship to evolution of catalytic activity remain uncertain. Here, we show that directed evolution of a computationally designed Kemp eliminase introduces dynamical changes that give rise to an activation heat capacity absent in the original design^3^. Extensive molecular dynamics simulations show that evolution results in the closure of solvent exposed loops and better packing of the active site with transition state stabilising residues. Remarkably, these changes give rise to a correlated dynamical network involving the transition state and large parts of the protein. This network tightens the transition state ensemble, which induces an activation heat capacity and thereby nonlinearity in the temperature dependence. Our results have implications for understanding enzyme evolution (e.g. in explaining the role of distal mutations and evolutionary tuning of dynamical responses) and suggest that integrating dynamics with design and evolution will accelerate the development of efficient novel enzymes.

---

Tailor-made enzymes promise to be transformative for the ‘green’ synthesis of pharmaceuticals and fine chemicals, and for achievement of a circular economy^4–6^. Increasingly, *de novo* computational design of catalytic residues and ligand binding pockets into protein scaffolds can afford incipient, yet generally modest, activity^7,8^. Such ‘designer enzymes’ can be significantly improved by directed evolution, which enhances activity by several orders of magnitude in the best cases^3,9–12^. This is now an effective way to develop new protein catalysts, but also demonstrates the limitations of current design protocols. Designed active sites often require fine-tuning by evolution to precisely position catalytic residues and tightly pack ligands^3,9–12^, increase electrostatic preorganization^13–16^, and reduce non-productive conformations^17–22^. What is less clear is how evolution acts on the overall protein scaffold, particularly its dynamics, to boost catalysis.

## Evolution of a negative activation heat capacity

The base-promoted Kemp elimination of benzisoxazoles is a valuable model for the catalysis of an elementary reaction^23,24^, and has become an exemplar for designed biological catalysis of a ‘non-natural’ reaction^3,11,12,25,26^. We previously subjected the computationally designed Kemp eliminase 1A53-2 to directed evolution (Fig. 1)^3^. Evolution boosted activity 10^4^-fold by introduction of six mutations during optimization of the first-shell residues only. The binding pocket of the resulting evolved variant 1A53-2.5 exhibits improved shape complementarity to the substrate and TS by virtue of several space-filling substitutions^3^. Introduction of the A157Y mutation furthermore enhances activity by restricting the conformational freedom and tuning the *pKa* of the catalytic base Glu178.

**Fig. 1.**
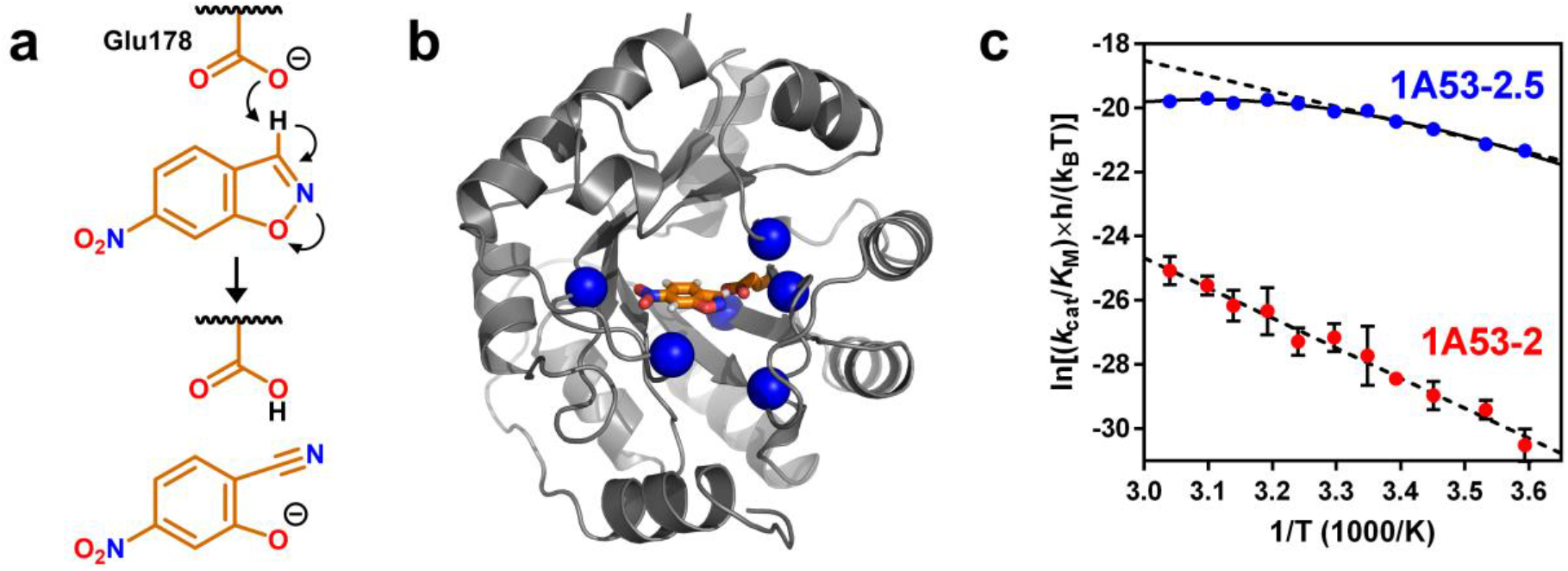
Activity gains during directed evolution of 1A32-2 coincided with the emergence of a negative 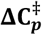. **a**, The general base Glu178 deprotonates 6-nitrobenzisoxazole. **b**, Six mutations (blue spheres) were introduced during active site optimization (ligand and base, orange carbons)^3^. **c**, The Eyring plot of 1A53-2 (red) is linear, whereas that of the evolved variant 1A53-2.5 (blue) is curved, signalling emergence of a negative 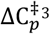. Dashed lines indicate a linear Eyring fit, solid lines show the macromolecular rate theory fit including 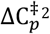.

Unexpectedly, activity gains in 1A53-2.5 coincided with the emergence of a curved temperature-dependence (Fig. 1c)^3^. This curvature is not due to protein unfolding or a change in mechanism. Rather, it can be attributed to temperature-dependent activation enthalpies and entropies due to an apparent activation heat capacity (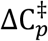, Equation 1). Such nonlinear temperature dependence is seen in many natural enzymes^1,2^, and is probably involved in thermoadaptation (Equation 2)^2^. Moreover, the pronounced 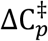 of some enzymes, including mesophiles, suggests a general relevance for enzyme catalysis^1–3^. A negative 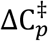 indicates that enthalpic fluctuations (A〈δH^2^〉^‡^) are reduced in the transition state ensemble compared to the enzyme-substrate complex^27^. Molecular dynamics (MD) simulations can probe this thermodynamic effect, and give 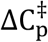 values in good agreement with experiment for two natural enzymes (Equation 3)^1^.

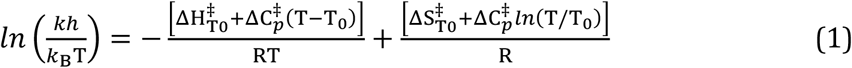

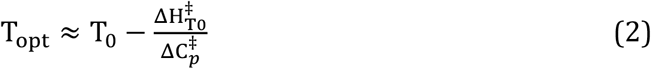

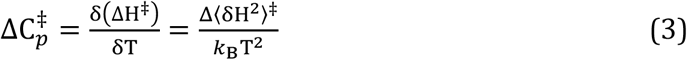

(*k*_B_: Boltzmann constant, T: Temperature, *h*: Planck constant, 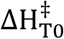 and 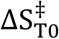 activation enthalpy and entropy at reference temperature T_0_, T_opt_: Temperature optimum)^2^.

Here, we show that evolution of 1A53-2 reshaped its dynamics, driving the emergence of a negative 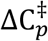. Strikingly, emergent correlated movements in the transition state ensemble of 1A53-2.5 span the protein and link the active site with the protein scaffold through a dynamical network. This network enhances catalytic preorganization and helps to explain the role of distal mutations that arise during further evolution.

## Energetic fluctuations

We performed 5 μs of MD simulations each for the designed and evolved variant in complex with a ground state (GS, Michaelis complex) and transition state model (TS, Extended Data Fig. 1). The designed and the evolved variants showed significantly different dynamical responses to the transition state. The energy distribution of 1A53-2.5 was notably narrower in the TS than in the GS. In contrast, no significant differences were observed for 1A53-2 (Extended Data Tab. 1+2, Extended Data Fig. 2a+b). The change in energetic fluctuations upon moving from GS to TS determines 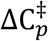, which we calculated from a moving average over the simulations using validated methods (Extended Data Fig. 2c)^1^. Rigidification of the TS ensemble in 1A53-2.5 results in a calculated negative 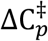 (−19.7 ± 3.1 kJ·mol^−1^K^−1^), whereas 1A53-2 shows no activation heat capacity (0.4 ± 3.2 kJ·mol^−1^K^−1^). This is consistent with the experimentally observed linear and nonlinear temperature dependence for the designed and evolved catalysts, respectively, and demonstrates the dynamical origin of 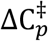.

The differences between the designed and evolved catalysts are significant. Errors for 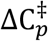 were calculated by cross-validation leaving out individual trajectories, and demonstrate the statistical significance of our calculations. Also, independent simulations using a transition state model with the transferring proton residing on the base (TS2, Extended Data Fig. 1) instead of the ligand gave similar results, supporting our findings (see Extended Data Tab. 1+2 and Extended Data Fig. 1,2,6-11 for TS2). We directly calculate 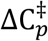 without fitting to experimental data from the difference in energetic fluctuations between GS and TS, after removal of bulk solvent. Our calculations predict a sizable negative 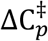 for the evolved protein, but the contributions of solvent likely reduce the magnitude of this effect substantially^1^ to yield the modest but still negative experimental value (−1.17 kJ·mol^−1^K^−1^). Also, slow loop movements, which cannot be exhaustively sampled on a reasonable timescale, may mix into the fluctuations at longer window sizes. In short, comparison of the designed and the evolved catalysts shows that the non-linear temperature dependence of the evolved variant is due to a 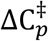 arising from dynamical differences between its GS and TS complex.

## Structural fluctuations

The decrease in energetic fluctuations associated with the reaction in 1A53-2.5 is accompanied by reduced structural fluctuations in the TS ensemble (Extended Data Fig. 7). In 1A53-2.5, several solvent-exposed loops covering the active site (residues 53-65, 84-92, and 181-192) become less mobile in the TS ensemble, as indicated by decreased root-mean square fluctuations (ΔRMSF) between GS and TS. Cluster analysis (Fig. 2a and Extended Data Fig. 8) and principal component analysis (Extended Data Fig. 9) of these loops shows that the scaffold interconverts between an open and a closed state. Notably, evolution increases the population of the closed state while also decreasing its radius of gyration (Extended Data Fig. 8), suggesting that pressure may modulate the fluctuations in 1A53-2.5 similar to temperature^28^.

**Fig. 2.**
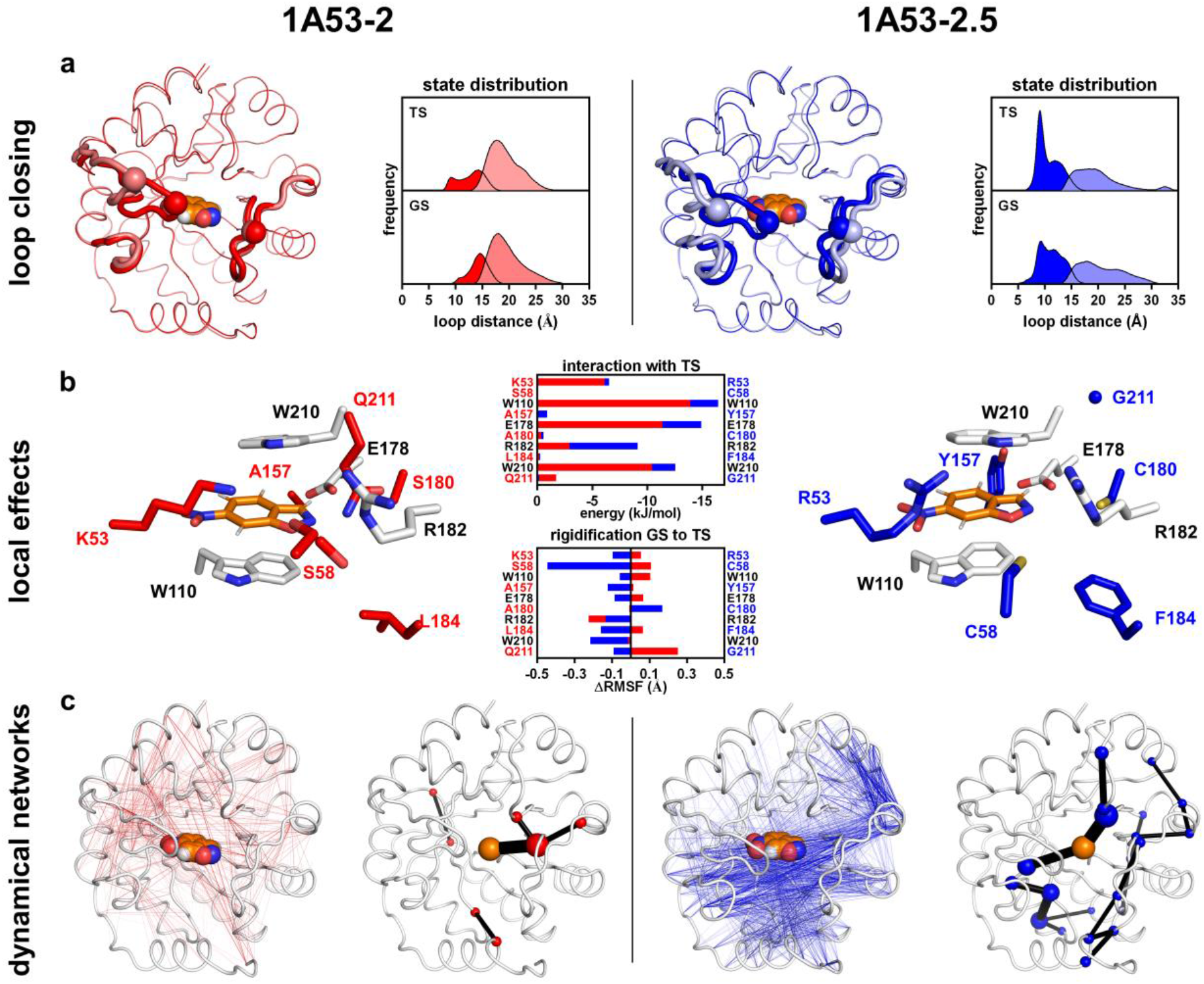
Global dynamics in the evolved enzyme are modulated by a local tightening of the active site associated reaction. **a,** Distance-based cluster analysis of the rigidifying loops (tubes) in 1A53-2 (red) and 1A53-2.5 (blue) reveals a conformational equilibrium between open (dark colors) and closed (light colors) states, as indicated e.g. by the Cα distance between residues 58 and 188 (spheres). The closed state becomes more populated as a result of evolution (see Extended Data Fig. 8+9). **b,** Evolution introduces six mutations at the active site (red/blue sticks, G211 as sphere) which enhance the interactions of Trp110, Glu178, Arg182 and Trp210 (grey sticks) with the TS in the closed state. As a result, the closed TS ensemble of 1A53-2.5 is more constrained than its GS ensemble, as reflected by reduced fluctuations (ΔRMSF) of many active site residues between the two states.

Loop closure affects active site preorganization in many enzymes, e.g. by desolvation and packing of ligands and catalytic residues^29–31^. In 1A53-2.5, the active site is tightly packed due to space-filling mutations such as A157Y and L184F^3^. Loop closure further enhances that packing as indicated by changes in the solvent accessible surface area of the ligand and base (Extended Data Fig. 11). This is partially achieved by expelling water from the active site, which is not observed for 1A53-2. Hydrogen bonding of Tyr157 with Glu178 in 1A53-2.5 additionally restricts the relative movement between the ligand and base. Notably, per-residue interaction energies with the TS indicate that the residues mutated during evolution do not directly provide additional TS stabilization (Fig. 2b). Instead, the mutations appear to strengthen interactions of the TS with other active site residues such as the catalytic base Glu178, Trp110 and Trp210 which sandwich the ligand, and Arg182 which stabilises the nascent oxyanion by long-range electrostatic interactions. In 1A53-2.5, this tightening is reflected by the ΔRMSF of several active site residues that are less mobile in the TS than in the GS. Evolution of these tight and ordered interactions enhanced active site preorganization, which we show here to be associated with the rigidification of the protein scaffold in the TS ensemble that causes the negative 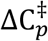.

## Evolution of a dynamical network

Dynamical correlations in the evolved TS ensemble connect local reorganization of the active site with the protein scaffold. The change in cross correlation between GS and TS was calculated for all backbone Cα atoms and the ligand, with the protein in the closed state (Fig. 3 and Extended Data Fig. 10). These correlations therefore reflect differences between the GS and TS, and do not involve conformational changes between the open and closed states. In the closed state of 1A53-2.5, large parts of the backbone move in a more correlated manner in the TS compared to the GS ensemble. Shortest pathway maps^19^ show that these increased correlations are communicated via neighbouring residues and indicate that evolution introduces a dynamical network that centres on the ligand and spans the protein (Fig. 3 and Extended Data Fig. 10). Only two mutations introduced during evolution are significantly involved in the network. Q211G introduces a flexible residue that potentially tunes the dynamic response of the scaffold, and L184F enhances packing by connecting neighbouring solvent-exposed loops. Instead of directly contributing to the network, the mutations apparently facilitate the global dynamical response to the TS by tightening its interactions with other active site residues (Trp110, Glu178, Arg182 and Trp220) as described above.

**Fig. 3.**
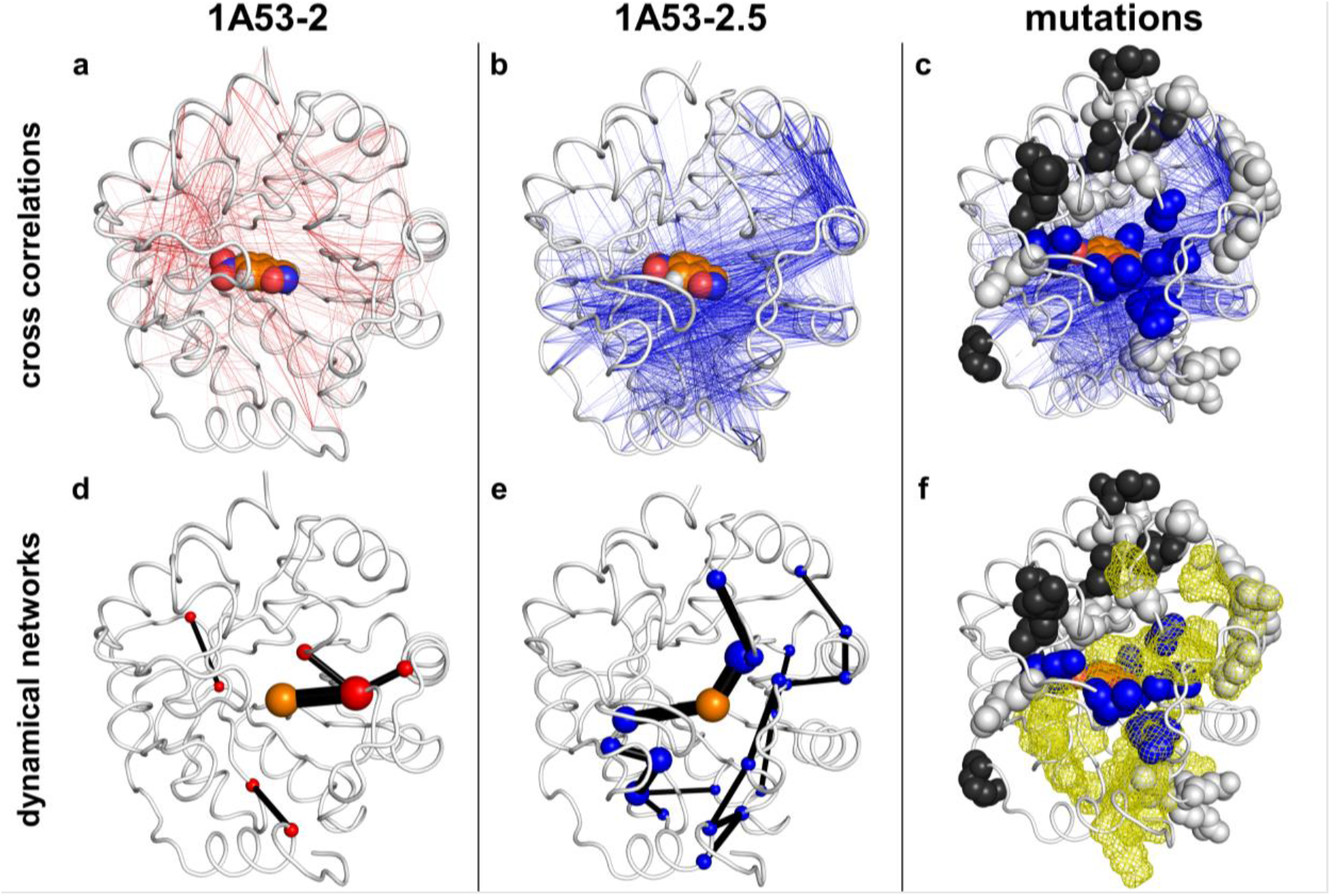
Directed evolution introduces an extended correlated network in the transition state ensemble. **a+b,** Evolution of 1A53-2 (red, left) to 1A53-2.5 (blue, middle) increases the correlated movements in the closed TS ensemble. Cross correlations that increase between GS and TS by ≥20% are indicated as lines on the structures (See Extended Data Fig. 10). **d+e,** Shortest pathway maps^19^ calculated from the increased cross correlations demonstrate evolution of a network centred on the chemical TS that spans the closed state of the protein. The sizes of the edges (black lines) and vertices (red/blue spheres: protein, orange spheres: ligand) indicate the strength of the network. **c+f,** The mutations in 1A53-2.5 (blue spheres), those found during further evolution using error-prone PCR (white spheres), and those in the final variant 1A53-2.9 (black spheres) cluster around the correlated movements **(c)** and network **(e**, yellow mesh) in 1A53-2.5, indicating fine-tuning of its dynamics by remote mutations.

1A53-2.5 was evolved by optimization of first shell residues only. During its further optimization by error-prone PCR, several distal mutations were identified that give an additional 3-fold activity improvement^3^. These mutations cluster around the increased cross correlations and the network in 1A53-2.5 (Fig. 3c+f), suggesting that these distal mutations modulate the dynamical network to enhance activity. The limited improvements achieved during these last rounds of evolution indicate that further fine-tuning would require many more mutations to reprogram the global dynamics encoded in the scaffold. While this may be challenging for laboratory evolution, Nature probably evolved similar networks to tailor activity, as indicated by phylogenetic analyses that revealed sectors of spatially proximal coevolving residues^32–34^. Notably, these sectors can also show correlated dynamics^35^, hinting at a catalytically relevant role of the underlying networks. Other allosteric effects apparently also rely on similar networks^36–38^, which may enhance preorganization, reduce non-productive conformations, and tune global scaffold flexibility for thermoadaptation^17–20,39^.

## Implications for catalysis and design

The co-emergence of catalytic power and 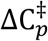 raises the intriguing question of the connection between transition state tightening (and the associated dynamical network) and catalysis^40^. While deciphering this cause-and-effect relationship will require further research, exquisite catalytic preorganization certainly requires ordering at the active site^15^. The evolved 1A53-2.5 achieves improved catalysis by increased TS stabilization, and its negative 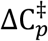 shows that the protein scaffold responds to the TS with significantly altered dynamics. We note that this does not imply that protein dynamics ‘drive’ the reaction on the timescale of the chemical bond making and breaking, which is supported by our QM/MM umbrella sampling simulations that reproduce the improvements achieved during evolution (Extended Data Fig. 3). Instead, effective TS stabilization involving exquisite active site preorganization is a function of the cooperative response of the whole protein scaffold. The transition from a flexible GS to a rigid TS, could thus lead to activity gains, albeit at the expense of introducing curved temperature-dependencies and therefore a loss of activity at elevated temperatures. 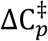 is apparently a signature of the superior catalytic efficiency achieved by the more ‘enzyme-like’ 1A53-2.5. Furthermore, 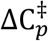 signals the ability of the scaffold to adapt to the chemical TS, which may relate to the evolvability of enzyme dynamics.

The dynamical network in 1A53-2.5 was evolved by introduction of only a few active-site mutations. Though computational design commonly targets active-site residues, the design of such networks will require accounting for dynamical changes associated with TS stabilization. Notably, our QM/MM simulations revealed that the improved catalytic environment that leads to these dynamical changes already takes effect at short timescales (<60 ps, Extended Data Fig. 3). Short, computationally efficient, MD simulations may thus reveal how mutations affect catalytic preorganization without the need to dissect the underlying dynamical networks. Furthermore, protein folds have highly conserved intrinsic dynamics^41^ that can be rapidly analysed for correlated sites^42^, with rigid anchor-points for catalytic residues^43^, and first-shell residues with high mobility and mutability^44^. Combining existing design algorithms with atomistically-detailed MD^1^ and rapid, approximate methods to model protein dynamics^42,45^ may thus help to attain tight communication with the scaffold’s dynamics and the exquisite precision required to achieve excellent TS complementarity and catalytic preorganization.

## Conclusion

Our atomistic molecular dynamics simulations and statistical thermodynamic analysis demonstrate how evolution of catalytic activity shapes protein dynamics. The catalytically superior 1A53-2.5 uniquely shows a correlated network and reduced fluctuations in the TS ensemble, which give rise to a negative 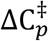. This finding suggests a connection between the cooperative response of the protein scaffold and TS stabilization at the active site. The results altogether imply that 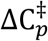 may be an indicator of successful catalytic preorganization and perhaps relates to evolvability. Integrating protein dynamics into design and evolution strategies may thus be essential for developing artificial enzymes that truly rival their natural counterparts.

## AUTHOR CONTRIBUTIONS

HAB, MWvdK and AJM devised the simulation and analysis. HAB performed the simulations and analysis. HAB, RA, DH, VLA, MWvdK, and AJM wrote the manuscript.

## ACKNOWLEDGMENTS

HAB and AJM thank EPSRC (EP/M013219/1 and EP/M022609/1) and with JRA BBSRC (BB/M000354/1) for funding. MWvdK is a BBSRC David Phillips Fellow (BB/M026280/1). VLA and AJM thank the Marsden Fund of New Zealand (16-UOW-027). VLA is a James Cook Research Fellow (Royal Society of New Zealand). DH thanks the Swiss National Science Foundation. This work was conducted using the computational facilities of the Advanced Computing Research Centre, University of Bristol. We thank Rory Crean and Silvia Osuna for help with and providing a script for performing the shortest-path analysis.

## REFERENCES

1 van der Kamp, M. W. et al. Dynamical origins of heat capacity changes in enzyme-catalysed reactions. Nat. Commun. 9, 1177 (2018).

2 Arcus, V. L. et al. On the temperature dependence of enzyme-catalyzed rates. Biochemistry 55, 1681–1688 (2016).

3 Bunzel, H. A. et al. Emergence of a negative activation heat capacity during evolution of a computationally designed enzyme. J. Am. Chem. Soc. 141, 11745–11748 (2019).

4 Arnold, F. H. Innovation by Evolution: Bringing New Chemistry to Life (Nobel Lecture). Angew. Chem. Int. Ed. 58, 14420–14426 (2019).

5 Bornscheuer, U. T. et al. Engineering the third wave of biocatalysis. Nature 485, 185–194 (2012).

6 Tournier, V. et al. An engineered PET depolymerase to break down and recycle plastic bottles. Nature 580, 216–219 (2020).

7 Kries, H., Blomberg, R. & Hilvert, D. De novo enzymes by computational design. Curr. Opin. Chem. Biol. 17, 221–228 (2013).

8 Kiss, G., Çelebi-Ölçüm, N., Moretti, R., Baker, D. & Houk, K. N. Computational enzyme design. Angew. Chem. Int. Ed. 52, 5700–5725 (2013).

9 Obexer, R. et al. Emergence of a catalytic tetrad during evolution of a highly active artificial aldolase. Nat. Chem. 9, 50–56 (2017).

10 Preiswerk, N. et al. Impact of scaffold rigidity on the design and evolution of an artificial Diels-Alderase. Proc. Natl. Acad. Sci. U. S. A. 111, 8013–8018 (2014).

11 Blomberg, R. et al. Precision is essential for efficient catalysis in an evolved Kemp eliminase. Nature 503, 418–421 (2013).

12 Khersonsky, O. et al. Bridging the gaps in design methodologies by evolutionary optimization of the stability and proficiency of designed Kemp eliminase KE59. Proc. Natl. Acad. Sci. U. S. A. 109, 10358–10363 (2012).

13 Fuxreiter, M. & Mones, L. The role of reorganization energy in rational enzyme design. Curr. Opin. Chem. Biol. 21, 34–41 (2014).

14 Jindal, G., Ramachandran, B., Bora, R. P. & Warshel, A. Exploring the development of ground-state destabilization and transition-state stabilization in two directed evolution paths of Kemp eliminases. ACS Catal. 7, 3301–3305 (2017).

15 Warshel, A. et al. Electrostatic basis for enzyme catalysis. Chem. Rev. 106, 3210–3235 (2006).

16 Bhowmick, A., Sharma, S. C. & Head-Gordon, T. The importance of the scaffold for de novo enzymes: A case study with Kemp eliminase. J. Am. Chem. Soc. 139, 5793–5800 (2017).

17 Hong, N. S. et al. The evolution of multiple active site configurations in a designed enzyme. Nat. Commun. 9, 3900 (2018).

18 Bhowmick, A., Sharma, S. C., Honma, H. & Head-Gordon, T. The role of side chain entropy and mutual information for improving the de novo design of Kemp eliminases KE07 and KE70. Phys. Chem. Chem. Phys. 18, 19386–19396 (2016).

19 Romero-Rivera, A., Garcia-Borras, M. & Osuna, S. Role of Conformational Dynamics in the Evolution of Retro-Aldolase Activity. ACS Catal. 7, 8524–8532 (2017).

20 Petrovic, D., Risso, V. A., Kamerlin, S. C. L. & Sanchez-Ruiz, J. M. Conformational dynamics and enzyme evolution. J. R. Soc. Interface 15 (2018).

21 Frushicheva, M. P. et al. Computer aided enzyme design and catalytic concepts. Curr. Opin. Chem. Biol. 21, 56–62 (2014).

22 Campbell, E. C. et al. Laboratory evolution of protein conformational dynamics. Curr. Opin. Struct. Biol. 50, 49–57 (2017).

23 Casey, M. L., Kemp, D. S., Paul, K. G. & Cox, D. D. Physical organic chemistry of benzisoxazoles. I. Mechanism of the base-catalyzed decomposition of benzisoxazoles. J. Org. Chem. 38, 2294–2301 (1973).

24 Kemp, D. S. & Casey, M. L. Physical organic chemistry of benzisoxazoles. II. Linearity of the Broensted free energy relation for the base-catalyzed decomposition of benzisoxazoles. J. Am. Chem. Soc. 95, 6670–6680 (1973).

25 Privett, H. K. et al. Iterative approach to computational enzyme design. Proc. Natl. Acad. Sci. U. S. A. 109, 3790–3795 (2012).

26 Röthlisberger, D. et al. Kemp elimination catalysts by computational enzyme design. Nature 453, 190–195 (2008).

27 Prabhu, N. V. & Sharp, K. A. Heat capacity in proteins. Annu. Rev. Phys. Chem. 56, 521–548 (2005).

28 Jones, H. B. L. et al. A complete thermodynamic analysis of enzyme turnover links the free energy landscape to enzyme catalysis. FEBS J. 284, 2829–2842 (2017).

29 Liao, Q. et al. Loop Motion in Triosephosphate Isomerase Is Not a Simple Open and Shut Case. J. Am. Chem. Soc. 140, 15889–15903 (2018).

30 van der Kamp, M. W., Chaudret, R. & Mulholland, A. J. QM/MM modelling of ketosteroid isomerase reactivity indicates that active site closure is integral to catalysis. FEBS J. 280, 3120–3131 (2013).

31 Malabanan, M. M., Amyes, T. L. & Richard, J. P. A role for flexible loops in enzyme catalysis. Curr. Opin. Struct. Biol. 20, 702–710 (2010).

32 Rivoire, O., Reynolds, K. A. & Ranganathan, R. Evolution-based functional decomposition of proteins. PLoS Comput. Biol. 12, e1004817 (2016).

33 Reynolds, K. A., McLaughlin, R. N. & Ranganathan, R. Hot spots for allosteric regulation on protein surfaces. Cell 147, 1564–1575 (2011).

34 Halabi, N., Rivoire, O., Leibler, S. & Ranganathan, R. Protein sectors: evolutionary units of three-dimensional structure. Cell 138, 774–786 (2009).

35 Lakhani, B., Thayer, K. M., Black, E. & Beveridge, D. L. Spectral analysis of molecular dynamics simulations on PDZ: MD sectors. J. Biomol. Struct. Dyn., 1–10 (2019).

36 Sethi, A., Eargle, J., Black, A. A. & Luthey-Schulten, Z. Dynamical networks in tRNA:protein complexes. Proc. Natl. Acad. Sci. U. S. A. 106, 6620–6625 (2009).

37 Rivalta, I. et al. Allosteric pathways in imidazole glycerol phosphate synthase. Proc. Natl. Acad. Sci. U. S. A. 109, E1428–1436 (2012).

38 Goodey, N. M. & Benkovic, S. J. Allosteric regulation and catalysis emerge via a common route. Nat. Chem. Biol. 4, 474–482 (2008).

39 Åqvist, J., Kazemi, M., Isaksen, G. V. & Brandsdal, B. O. Entropy and enzyme catalysis. Acc. Chem. Res. 50, 199–207 (2017).

40 Williams, D. H., Stephens, E., O’Brien, D. P. & Zhou, M. Understanding Noncovalent Interactions: Ligand Binding Energy and Catalytic Efficiency from Ligand-Induced Reductions in Motion within Receptors and Enzymes. Angew. Chem. Int. Ed. 43, 6596–6616 (2004).

41 Zhang, S., Li, H., Krieger, J. M. & Bahar, I. Shared Signature Dynamics Tempered by Local Fluctuations Enables Fold Adaptability and Specificity. Mol. Biol. Evol. 36, 2053–2068 (2019).

42 Tiwari, S. P. & Reuter, N. Conservation of intrinsic dynamics in proteins-what have computational models taught us? Curr. Opin. Struct. Biol. 50, 75–81 (2018).

43 Fuglebakk, E., Echave, J. & Reuter, N. Measuring and comparing structural fluctuation patterns in large protein datasets. Bioinformatics 28, 2431–2440 (2012).

44 Echave, J., Spielman, S. J. & Wilke, C. O. Causes of evolutionary rate variation among protein sites. Nat. Rev. Genet. 17, 109–121 (2016).

45 Wells, S. A., van der Kamp, M. W., McGeagh, J. D. & Mulholland, A. J. Structure and Function in Homodimeric Enzymes: Simulations of Cooperative and Independent Functional Motions. PloS one 10, e0133372 (2015).

46 Na, J., Houk, K. N. & Hilvert, D. Transition state of the base-promoted ring-opening of isoxazoles. Theoretical prediction of catalytic functionalities and design of haptens for antibody production. J. Am. Chem. Soc. 118, 6462–6471 (1996).

47 Alexandrova, A. N., Röthlisberger, D., Baker, D. & Jorgensen, W. L. Catalytic Mechanism and Performance of Computationally Designed Enzymes for Kemp Elimination. J. Am. Chem. Soc. 130, 15907–15915 (2008).

48 Swiderek, K., Tunon, I., Moliner, V. & Bertran, J. Revealing the origin of the efficiency of the de novo designed Kemp eliminase HG-3.17 by comparison with the former developed HG-3. Chemistry 23, 7582–7589 (2017).

