## Extended Data and Figures for "Evolution of dynamical networks enhances catalysis in a designer enzyme"

Extended Tables and Figures for

**Extended Data Table 1 | Variances and** $\boldsymbol{\Delta C}_{\boldsymbol{p}}^{\boldsymbol{\ddagger}}$ **of 1A53-2.^a^**

| Run | Moving Window (ns) | | | | | | | | | | | | | | | |
| --- | --- | --- | --- | --- | --- | --- | --- | --- | --- | --- | --- | --- | --- | --- | --- | --- |
|  | 5 | 10 | 15 | 20 | 25 | 30 | 35 | 40 | 45 | 50 | 55 | 60 | 65 | 70 | 75 | 80 |
| 1A53-2 : GS : Variance (kJ^2^∙mol^-2^) | | | | | | | | | | | | | | | | |
| 1 | 50910 | 57655 | 61152 | 62914 | 63994 | 64789 | 65588 | 66334 | 67011 | 67640 | 68196 | 68760 | 69337 | 69846 | 70341 | 70810 |
| 2 | 51915 | 59229 | 63796 | 66724 | 69029 | 71070 | 72799 | 74376 | 75848 | 77282 | 78699 | 80111 | 81436 | 82787 | 84152 | 85453 |
| 3 | 52809 | 61675 | 66191 | 69640 | 71983 | 73482 | 74545 | 75547 | 76371 | 76951 | 77359 | 77704 | 78039 | 78427 | 78796 | 79030 |
| 4 | 51217 | 59484 | 64315 | 67783 | 70043 | 71676 | 73080 | 74268 | 75405 | 76501 | 77454 | 78334 | 79206 | 80120 | 81031 | 81884 |
| 5 | 47598 | 52665 | 55491 | 57261 | 58389 | 59334 | 60130 | 60918 | 61544 | 62040 | 62443 | 62742 | 63040 | 63376 | 63749 | 64145 |
| 6 | 48169 | 56226 | 61256 | 64799 | 67675 | 70269 | 72691 | 74894 | 76905 | 78809 | 80624 | 82394 | 84116 | 85806 | 87437 | 88954 |
| 7 | 50943 | 57331 | 60427 | 62257 | 63658 | 64896 | 65835 | 66591 | 67241 | 67817 | 68357 | 68805 | 69217 | 69620 | 69958 | 70208 |
| 8 | 51756 | 58140 | 61448 | 64017 | 66066 | 67834 | 69472 | 70874 | 72066 | 73082 | 73971 | 74723 | 75405 | 76004 | 76484 | 76960 |
| 9 | 48671 | 56364 | 60778 | 63831 | 66315 | 68512 | 70554 | 72185 | 73412 | 74388 | 75296 | 76192 | 77043 | 77861 | 78598 | 79310 |
| 10 | 51151 | 58329 | 62287 | 64308 | 65603 | 66617 | 67582 | 68490 | 69353 | 70080 | 70729 | 71205 | 71483 | 71775 | 72071 | 72243 |
| Average | 50514 | 57710 | 61714 | 64353 | 66275 | 67848 | 69228 | 70448 | 71516 | 72459 | 73313 | 74097 | 74832 | 75562 | 76262 | 76900 |
| Std. Dev. | 1745 | 2389 | 2854 | 3378 | 3816 | 4152 | 4471 | 4778 | 5089 | 5408 | 5734 | 6097 | 6467 | 6851 | 7232 | 7600 |
| 1A53-2 : TS : Variance (kJ^2^∙mol^-2^) | | | | | | | | | | | | | | | | |
| 1 | 50084 | 57964 | 63380 | 67196 | 70074 | 72196 | 73766 | 74863 | 75488 | 75918 | 76332 | 76903 | 77631 | 78431 | 79175 | 79809 |
| 2 | 49238 | 57735 | 61656 | 63633 | 64893 | 65948 | 66928 | 67801 | 68644 | 69389 | 70054 | 70597 | 70970 | 71261 | 71403 | 71510 |
| 3 | 48644 | 53988 | 56753 | 58618 | 60137 | 61462 | 62607 | 63649 | 64604 | 65381 | 65962 | 66449 | 66831 | 67069 | 67201 | 67311 |
| 4 | 49172 | 57299 | 62168 | 65496 | 68197 | 70415 | 72204 | 73685 | 74883 | 75841 | 76479 | 76823 | 77037 | 77178 | 77305 | 77473 |
| 5 | 47174 | 54805 | 58768 | 61587 | 63754 | 65243 | 66231 | 67077 | 67773 | 68414 | 68920 | 69320 | 69616 | 69786 | 69917 | 70065 |
| 6 | 50956 | 58731 | 63370 | 66687 | 69442 | 71728 | 73633 | 75243 | 76295 | 77047 | 77541 | 77982 | 78367 | 78686 | 78998 | 79306 |
| 7 | 49302 | 54705 | 57171 | 58658 | 59751 | 60809 | 61708 | 62468 | 63210 | 63894 | 64564 | 65179 | 65732 | 66260 | 66749 | 67258 |
| 8 | 49962 | 58007 | 62890 | 66632 | 69400 | 71252 | 72387 | 73332 | 74481 | 75817 | 77171 | 78459 | 79563 | 80605 | 81529 | 82165 |
| 9 | 50341 | 57941 | 63019 | 66479 | 68663 | 69968 | 70943 | 72078 | 73427 | 74763 | 75973 | 77099 | 78280 | 79433 | 80557 | 81555 |
| 10 | 51625 | 59497 | 64449 | 67757 | 70363 | 72481 | 74276 | 75958 | 77454 | 78758 | 80000 | 81190 | 82362 | 83534 | 84667 | 85761 |
| Average | 49650 | 57067 | 61362 | 64274 | 66467 | 68150 | 69468 | 70615 | 71626 | 72522 | 73300 | 74000 | 74639 | 75224 | 75750 | 76221 |
| Std. Dev. | 1249 | 1880 | 2768 | 3489 | 4062 | 4446 | 4730 | 4970 | 5127 | 5270 | 5418 | 5595 | 5819 | 6091 | 6392 | 6655 |
| Av. TS1-GS | -864 | -643 | -352 | -79 | 192 | 302 | 241 | 168 | 110 | 63 | -13 | -97 | -193 | -338 | -512 | -679 |
| 1A53-2 : TS - GS : ∆C^‡^*_p_* (kJ∙mol^-1^) | | | | | | | | | | | | | | | | |
| **∆C^‡^_p_** | **-4.9** | **-3.6** | **-2.0** | **-0.4** | **1.1** | **1.7** | **1.4** | **0.9** | **0.6** | **0.4** | **-0.1** | **-0.6** | **-1.1** | **-1.9** | **-2.9** | **-3.8** |
| **∆∆C^‡^_p_** | **0.9** | **1.3** | **1.7** | **2.1** | **2.4** | **2.6** | **2.7** | **2.9** | **3.1** | **3.2** | **3.3** | **3.5** | **3.7** | **3.9** | **4.1** | **4.3** |
| 1A53-2 : TS2 : Variance (kJ^2^∙mol^-2^) | | | | | | | | | | | | | | | | |
| 1 | 52987 | 62195 | 67280 | 70491 | 72911 | 74978 | 76726 | 78175 | 79594 | 80946 | 82404 | 83960 | 85533 | 87091 | 88556 | 89867 |
| 2 | 46339 | 52620 | 56149 | 58257 | 60013 | 61683 | 63181 | 64462 | 65682 | 66821 | 67704 | 68508 | 69324 | 70166 | 70982 | 71797 |
| 3 | 47702 | 55226 | 59193 | 61710 | 63467 | 64795 | 65903 | 66979 | 68006 | 69000 | 70087 | 71101 | 72087 | 73129 | 74202 | 75301 |
| 4 | 55094 | 61686 | 65016 | 67325 | 69009 | 70109 | 70902 | 71400 | 71857 | 72362 | 72864 | 73402 | 73953 | 74338 | 74735 | 75121 |
| 5 | 50878 | 58694 | 62870 | 65436 | 67364 | 69019 | 70596 | 72041 | 73137 | 74209 | 75350 | 76366 | 77314 | 78160 | 78831 | 79433 |
| 6 | 48514 | 54373 | 57097 | 58684 | 60221 | 61570 | 62820 | 63999 | 65077 | 66057 | 66977 | 67888 | 68769 | 69577 | 70398 | 71269 |
| 7 | 48849 | 55733 | 59649 | 62575 | 64627 | 66032 | 66940 | 67763 | 68577 | 69283 | 69817 | 70275 | 70728 | 71194 | 71672 | 72109 |
| 8 | 51307 | 59505 | 63619 | 67044 | 69891 | 72138 | 73326 | 74230 | 74987 | 75794 | 76619 | 77409 | 78238 | 79123 | 79833 | 80379 |
| 9 | 47717 | 55057 | 60650 | 65012 | 68520 | 71692 | 74637 | 77210 | 79571 | 82008 | 84460 | 86856 | 89070 | 91171 | 93188 | 95191 |
| 10 | 51667 | 58007 | 61288 | 64023 | 66464 | 68554 | 70378 | 72021 | 73498 | 74834 | 76103 | 77176 | 78044 | 78826 | 79463 | 79985 |
| Average | 50105 | 57310 | 61281 | 64056 | 66249 | 68057 | 69541 | 70828 | 71999 | 73132 | 74239 | 75294 | 76306 | 77277 | 78186 | 79045 |
| Std. Dev. | 2737 | 3210 | 3483 | 3859 | 4181 | 4480 | 4728 | 4954 | 5196 | 5511 | 5922 | 6370 | 6797 | 7216 | 7602 | 7966 |
| Av. TS2-GS | -408 | -400 | -433 | -298 | -27 | 209 | 313 | 380 | 483 | 673 | 926 | 1197 | 1474 | 1715 | 1924 | 2146 |
| 1A53-2 : TS2 - GS : ∆C^‡^*_p_* (kJ∙mol^-1^) | | | | | | | | | | | | | | | | |
| **∆C^‡^_p_** | **-2.3** | **-2.3** | **-2.5** | **-1.7** | **-0.2** | **1.2** | **1.8** | **2.2** | **2.7** | **3.8** | **5.2** | **6.8** | **8.3** | **9.7** | **10.9** | **12.2** |
| **∆∆C^‡^_p_** | **1.4** | **1.7** | **1.9** | **2.2** | **2.4** | **2.6** | **2.7** | **2.9** | **3.1** | **3.3** | **3.5** | **3.7** | **4.0** | **4.2** | **4.4** | **4.7** |

^a^ Variances and heat capacities calculate for different moving average windows, based on complexes comprising the protein, ligand, and ten water molecules closest to Glu178.

**Extended Data Table 2 | Variances and** $\boldsymbol{\Delta C}_{\boldsymbol{p}}^{\boldsymbol{\ddagger}}$ **for 1A53-2.5.^a^**

| Run | Moving Window (ns) | | | | | | | | | | | | | | | |
| --- | --- | --- | --- | --- | --- | --- | --- | --- | --- | --- | --- | --- | --- | --- | --- | --- |
|  | 5 | 10 | 15 | 20 | 25 | 30 | 35 | 40 | 45 | 50 | 55 | 60 | 65 | 70 | 75 | 80 |
| 1A53-2.5 : GS : Variance (kJ^2^∙mol^-2^) | | | | | | | | | | | | | | | | |
| 1 | 47802 | 54132 | 57931 | 60672 | 63142 | 65404 | 67296 | 68901 | 70238 | 71456 | 72619 | 73668 | 74624 | 75549 | 76490 | 77428 |
| 2 | 50645 | 59588 | 64984 | 68618 | 71527 | 73730 | 75063 | 75872 | 76323 | 76640 | 76893 | 77107 | 77268 | 77408 | 77639 | 77955 |
| 3 | 52923 | 61034 | 65056 | 67862 | 70284 | 72278 | 73872 | 75245 | 76515 | 77697 | 78814 | 79753 | 80639 | 81431 | 82198 | 83008 |
| 4 | 47297 | 53118 | 56709 | 59841 | 62750 | 65254 | 67074 | 68173 | 68930 | 69396 | 69735 | 70095 | 70497 | 70960 | 71497 | 72028 |
| 5 | 47567 | 54338 | 58478 | 61619 | 64406 | 66912 | 69196 | 71199 | 73108 | 74890 | 76560 | 78118 | 79545 | 80833 | 81838 | 82725 |
| 6 | 47155 | 53959 | 57291 | 59536 | 61334 | 62803 | 64099 | 65243 | 66262 | 67152 | 68025 | 68840 | 69646 | 70472 | 71289 | 72118 |
| 7 | 52035 | 61936 | 67582 | 70893 | 73190 | 75231 | 77096 | 78767 | 80034 | 81097 | 82128 | 83043 | 83875 | 84725 | 85579 | 86412 |
| 8 | 48048 | 55662 | 59831 | 62331 | 64164 | 65420 | 66386 | 67068 | 67686 | 68165 | 68501 | 68867 | 69173 | 69445 | 69661 | 69848 |
| 9 | 48682 | 53949 | 57399 | 60023 | 62286 | 64049 | 65345 | 66346 | 67058 | 67630 | 68199 | 68794 | 69408 | 70059 | 70738 | 71397 |
| 10 | 44837 | 50595 | 54482 | 57614 | 60012 | 61913 | 63491 | 65000 | 66431 | 67656 | 68670 | 69393 | 69967 | 70333 | 70547 | 70685 |
| Average | 48699 | 55831 | 59974 | 62901 | 65309 | 67299 | 68892 | 70182 | 71259 | 72178 | 73014 | 73768 | 74464 | 75121 | 75748 | 76360 |
| Std. Dev. | 2462 | 3733 | 4345 | 4534 | 4618 | 4718 | 4797 | 4877 | 4931 | 5033 | 5191 | 5350 | 5509 | 5674 | 5837 | 6013 |
| 1A53-2.5 : TS : Variance (kJ^2^∙mol^-2^) | | | | | | | | | | | | | | | | |
| 1 | 47671 | 54854 | 59147 | 62004 | 63978 | 65377 | 66510 | 67460 | 68436 | 69344 | 70169 | 71005 | 71878 | 72735 | 73479 | 74182 |
| 2 | 45799 | 50643 | 54030 | 56473 | 58183 | 59542 | 60711 | 61803 | 62795 | 63756 | 64698 | 65410 | 65980 | 66512 | 66971 | 67411 |
| 3 | 49222 | 56868 | 60907 | 62935 | 64275 | 65530 | 66740 | 67967 | 69261 | 70554 | 71782 | 72972 | 74079 | 75170 | 76238 | 77232 |
| 4 | 45963 | 51755 | 54561 | 56497 | 58020 | 59216 | 60156 | 60874 | 61489 | 62053 | 62583 | 63081 | 63542 | 64043 | 64600 | 65190 |
| 5 | 51568 | 58750 | 62496 | 64978 | 67023 | 68630 | 69914 | 70983 | 71988 | 72829 | 73580 | 74207 | 74758 | 75270 | 75736 | 76065 |
| 6 | 45092 | 50995 | 53978 | 55661 | 56777 | 57789 | 58685 | 59307 | 59801 | 60313 | 60786 | 61248 | 61624 | 61960 | 62266 | 62533 |
| 7 | 50525 | 58326 | 63271 | 66639 | 68959 | 70754 | 72503 | 74333 | 76104 | 77725 | 79087 | 80221 | 81280 | 82241 | 83174 | 84049 |
| 8 | 48975 | 55820 | 59755 | 62395 | 64320 | 65853 | 66986 | 68047 | 69105 | 70182 | 71217 | 72236 | 73262 | 74113 | 74705 | 75290 |
| 9 | 50011 | 56785 | 60605 | 63058 | 64977 | 66576 | 67988 | 69291 | 70371 | 71283 | 72049 | 72724 | 73038 | 73174 | 73306 | 73486 |
| 10 | 50150 | 56308 | 60235 | 62661 | 64283 | 65700 | 66813 | 67581 | 68252 | 68903 | 69551 | 70278 | 71067 | 71860 | 72609 | 73284 |
| Average | 48498 | 55110 | 58898 | 61330 | 63080 | 64497 | 65701 | 66765 | 67760 | 68694 | 69550 | 70338 | 71051 | 71708 | 72309 | 72872 |
| Std. Dev. | 2244 | 2975 | 3469 | 3793 | 4058 | 4257 | 4456 | 4722 | 5015 | 5273 | 5482 | 5666 | 5850 | 6013 | 6166 | 6307 |
| Av. TS1-GS | -201 | -721 | -1076 | -1571 | -2230 | -2803 | -3191 | -3417 | -3498 | -3484 | -3464 | -3430 | -3413 | -3414 | -3439 | -3488 |
| 1A53-2.5 : TS - GS : ∆C^‡^*_p_* (kJ∙mol^-1^) | | | | | | | | | | | | | | | | |
| **∆C^‡^_p_** | **-1.1** | **-4.1** | **-6.1** | **-8.9** | **-12.6** | **-15.9** | **-18.1** | **-19.4** | **-19.8** | **-19.7** | **-19.6** | **-19.4** | **-19.3** | **-19.3** | **-19.5** | **-19.8** |
| **∆∆C^‡^_p_** | **1.4** | **2.0** | **2.3** | **2.5** | **2.6** | **2.7** | **2.8** | **2.9** | **3.0** | **3.1** | **3.2** | **3.3** | **3.4** | **3.5** | **3.6** | **3.7** |
| 1A53-2.5 : TS2 : Variance (kJ^2^∙mol^-2^) | | | | | | | | | | | | | | | | |
| 1 | 51150 | 59880 | 65304 | 69075 | 71945 | 74284 | 76070 | 77533 | 78675 | 79691 | 80671 | 81607 | 82468 | 83212 | 83746 | 84115 |
| 2 | 46482 | 52393 | 55830 | 58598 | 60909 | 62824 | 64516 | 65951 | 67211 | 68303 | 69225 | 69962 | 70473 | 70941 | 71404 | 71850 |
| 3 | 51810 | 58509 | 61906 | 64301 | 66224 | 67811 | 69272 | 70713 | 71928 | 72959 | 73630 | 74112 | 74587 | 75025 | 75465 | 75930 |
| 4 | 50457 | 56486 | 59927 | 62276 | 64074 | 65374 | 66190 | 66963 | 67714 | 68443 | 69175 | 69835 | 70483 | 71163 | 71837 | 72451 |
| 5 | 48669 | 55851 | 59778 | 61458 | 62056 | 62635 | 63360 | 64037 | 64598 | 65148 | 65636 | 66104 | 66536 | 66958 | 67354 | 67717 |
| 6 | 48182 | 54295 | 57205 | 58517 | 59346 | 60021 | 60628 | 61154 | 61594 | 61999 | 62377 | 62680 | 63007 | 63382 | 63774 | 64131 |
| 7 | 50854 | 57652 | 61154 | 63882 | 65759 | 66959 | 67693 | 68126 | 68494 | 68914 | 69340 | 69628 | 69948 | 70326 | 70691 | 71051 |
| 8 | 47770 | 54890 | 59516 | 63103 | 65714 | 67850 | 69641 | 71379 | 73198 | 75105 | 76984 | 78730 | 80393 | 81962 | 83363 | 84581 |
| 9 | 47872 | 54204 | 57690 | 60089 | 62019 | 63733 | 65140 | 66262 | 67071 | 67872 | 68708 | 69513 | 70277 | 71032 | 71828 | 72662 |
| 10 | 49319 | 56343 | 60446 | 62831 | 64404 | 65691 | 66945 | 68143 | 69132 | 70037 | 70768 | 71295 | 71739 | 72074 | 72324 | 72546 |
| Average | 49256 | 56050 | 59876 | 62413 | 64245 | 65718 | 66945 | 68026 | 68962 | 69847 | 70651 | 71347 | 71991 | 72608 | 73179 | 73703 |
| Std. Dev. | 1746 | 2231 | 2666 | 3106 | 3537 | 3919 | 4203 | 4479 | 4748 | 5023 | 5299 | 5583 | 5863 | 6107 | 6295 | 6443 |
| Av. TS2-GS | 558 | 219 | -99 | -488 | -1064 | -1581 | -1946 | -2156 | -2297 | -2331 | -2363 | -2421 | -2473 | -2514 | -2569 | -2657 |
| 1A53-2.5 : TS2 - GS : ∆C^‡^*_p_* (kJ∙mol^-1^) | | | | | | | | | | | | | | | | |
| **∆C^‡^_p_** | **3.2** | **1.2** | **-0.6** | **-2.8** | **-6.0** | **-9.0** | **-11.0** | **-12.2** | **-13.0** | **-13.2** | **-13.4** | **-13.7** | **-14.0** | **-14.3** | **-14.6** | **-15.1** |
| **∆∆C^‡^_p_** | **1.3** | **1.8** | **2.2** | **2.3** | **2.5** | **2.6** | **2.7** | **2.8** | **2.9** | **3.0** | **3.1** | **3.3** | **3.4** | **3.5** | **3.6** | **3.7** |

^a^ Variances and heat capacities calculate for different moving average windows, based on complexes comprising the protein, ligand, and ten water molecules closest to Glu178.

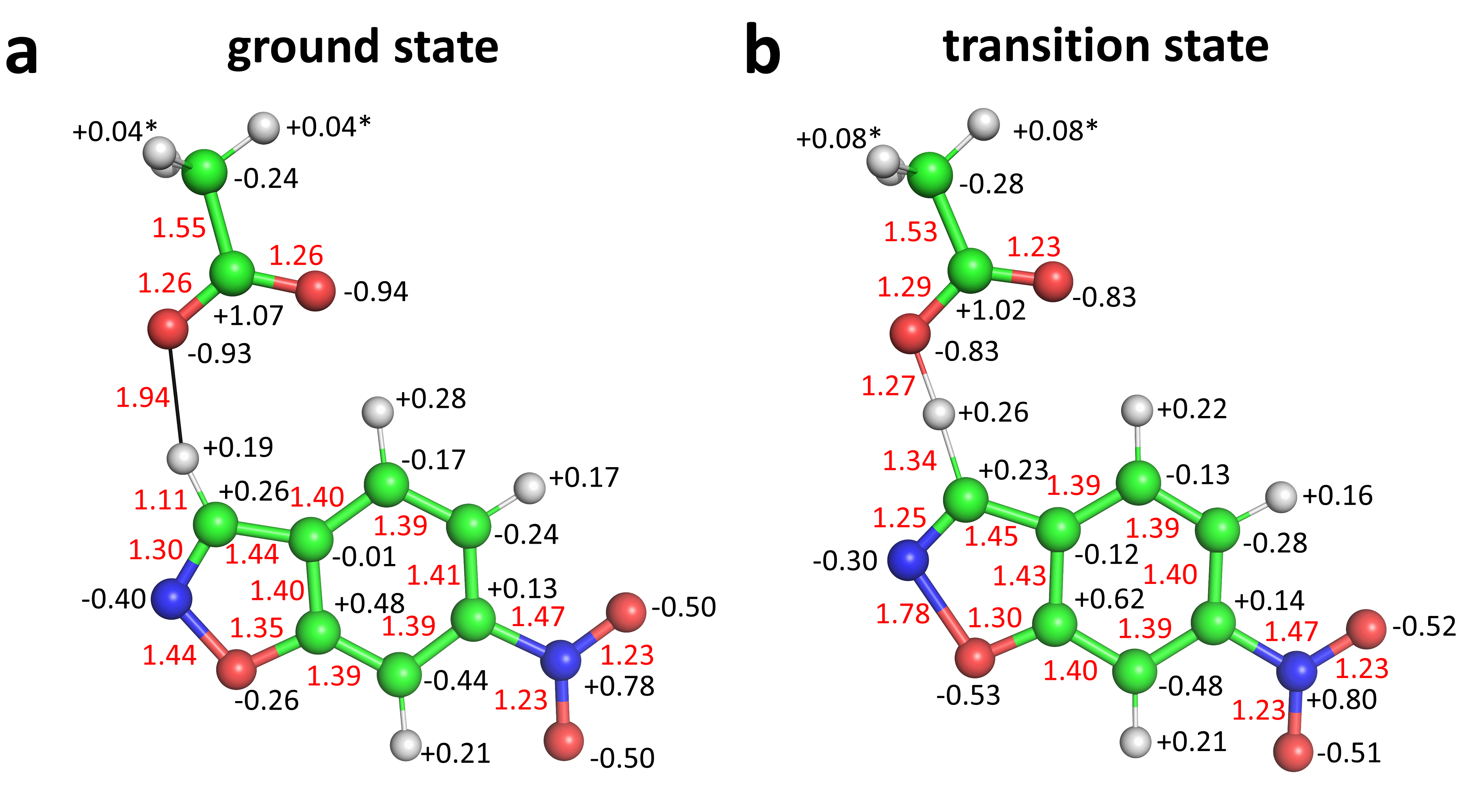

**Extended Data Fig. 1** **| MD parameters of the ground state and transition state model.** Distances (red) and charges (black) are based on gas-phase calculations with 6-nitrobenzisoxazole and acetate. To prevent artificial rigidification of the transition state by introduction of a covalent bond between the ligand and catalytic residue,^1^ this state was simulated with the transferring protein residing either on the ligand (TS) or base (TS2). Both transition state models provided similar results, supporting the significance of our findings. ***,** Because Glu178 has only two γ-hydrogens, their charge is adjusted to the given value according to the corresponding three hydrogens in acetate.

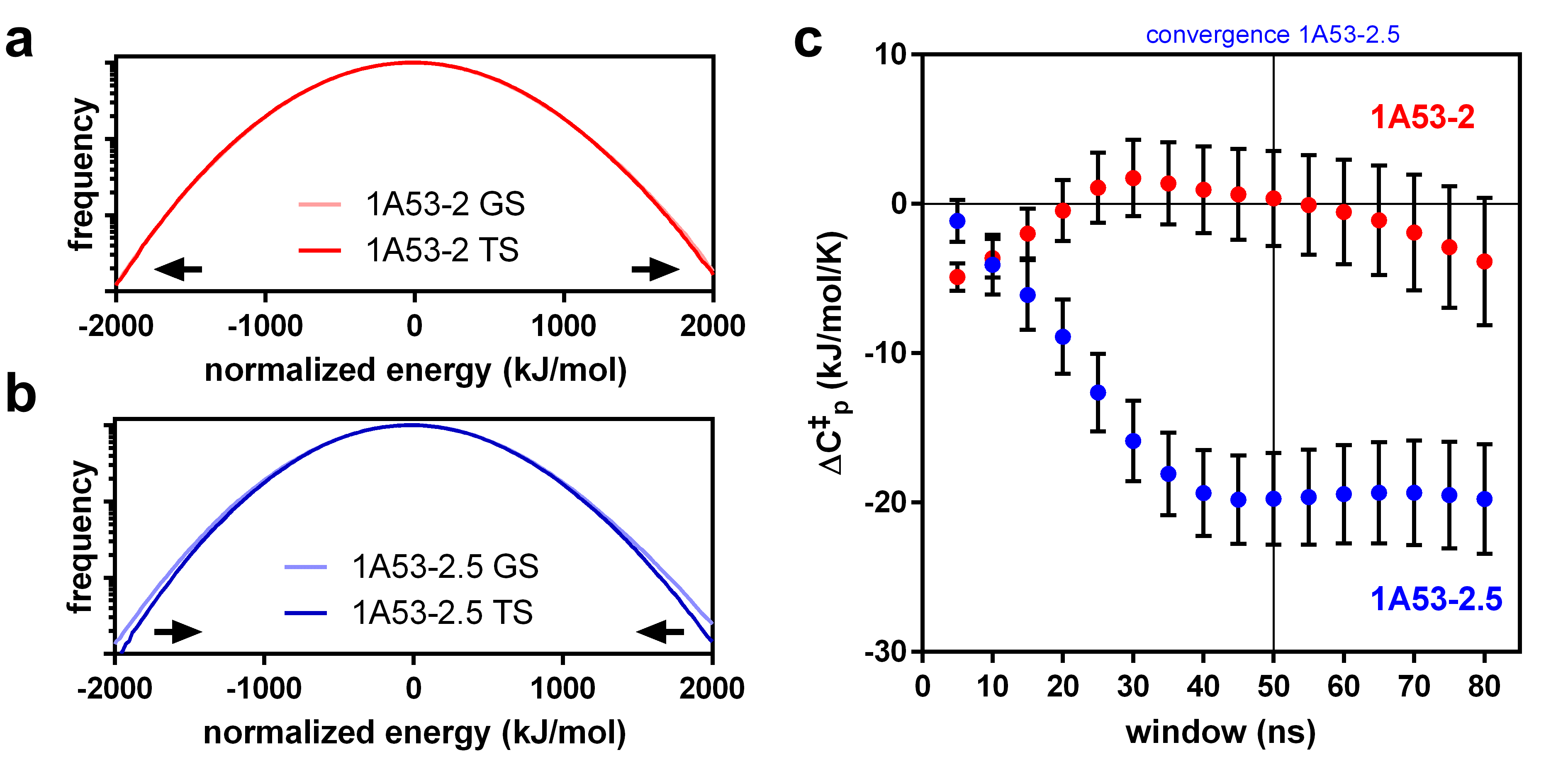

**Extended Data Fig. 2 | Changes in energetic fluctuations give rise to an activation heat capacity** $\boldsymbol{\Delta C}_{\boldsymbol{p}}^{\boldsymbol{\ddagger}}$ **in the evolved catalyst.** **a,** The energy distributions for 1A53-2 (TS: dark red; GS: light red) are almost identical, indicating a ${\Delta C}_{p}^{\ddagger}$ close to zero. Energetic fluctuations were calculated after removal of all but 10 water molecules, though removing these only slightly affected ${\Delta C}_{p}^{\ddagger}$ (Extended Data Fig. 6). **b,** The distribution for 1A53-2.5 is narrower for the TS (dark blue) than for the GS (light blue) complex, resulting in a negative ${\Delta C}_{p}^{\ddagger}$. Distributions in **a** and **b** are normalized to a 50 ns moving average and plotted against a log scale to emphasize changes in variance. **c,** ${\Delta C}_{p}^{\ddagger}$ was calculated based on the difference in variance between the TS and GS ensemble for moving windows of various sizes, and converges after 50 ns to negative values for 1A53-2.5 (blue) and to zero for 1A53-2 (red). Error bars were obtained by leave-one-out cross-validation.

|  | **a** **PM6/CHARMM36** | **b** **AM1/CHARMM36** | **c** **PM6/AMBERff99SB** |
| --- | --- | --- | --- |
| **1A53-2** | 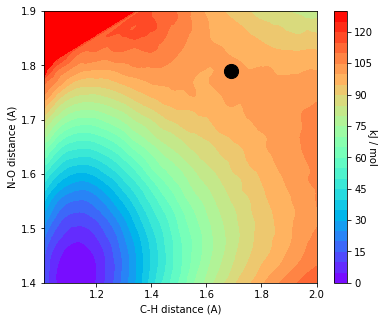 | 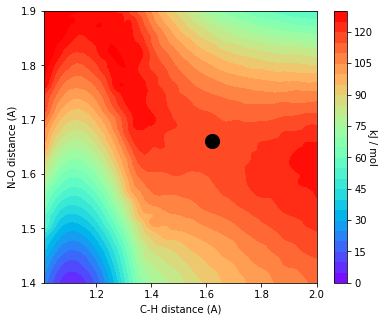 | 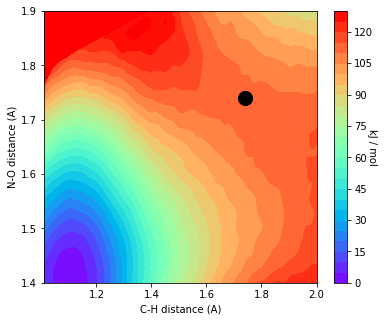 |
| **1A53-2.5** | 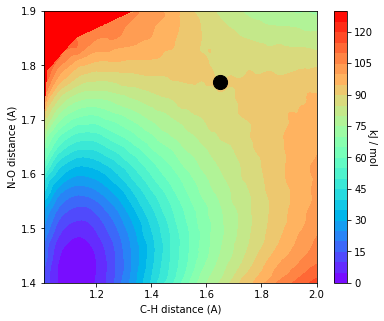 | 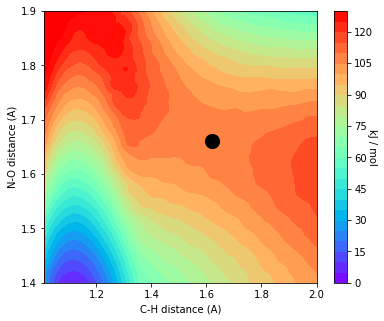 | 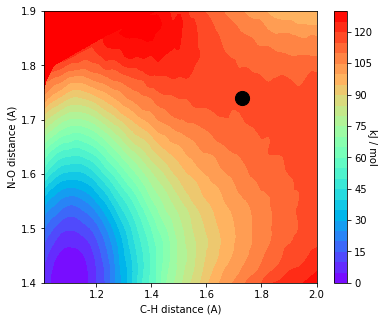 |

**Extended Data Fig. 3 | QM/MM modelling of the Kemp eliminases.** Free energy surfaces of 1A53-2 (top) and 1A53-2.5 (bottom) at the **a,** PM6/CHARMM36, **b,** AM1/CHARMM36 and the **c,** PM6/AMBERff99SB level of theory. The ligand as well as the carboxylate of Glu178 were part of the QM region during umbrella sampling of the C-H and N-O bonds. The transition state (black dot) reproduces the asynchronous Kemp elimination with C-H cleavage preceding N-O cleavage observed for similar systems.^46-48^ PM6/CHARMM36 gave activation energies of 101 kJ∙mol^‑1^ and 90 kJ∙mol^‑1^, which reproduce the improvements achieved during evolution (86 and 68 kJ∙mol^‑1^ for *k*_cat_). Similarly, simulations with AM1/CHARMM36 preserved that trend, but gave higher energies (117 and 105 kJ∙mol^‑1^). In contrast, simulations with PM6/AMBERff99SB resulted in an inverse trend for the activation energies (111 and 116 kJ∙mol^‑1^). This observation for AMBERff99SB mimics the classical MD, in which the ligand was expelled from the active site within 100 ns.

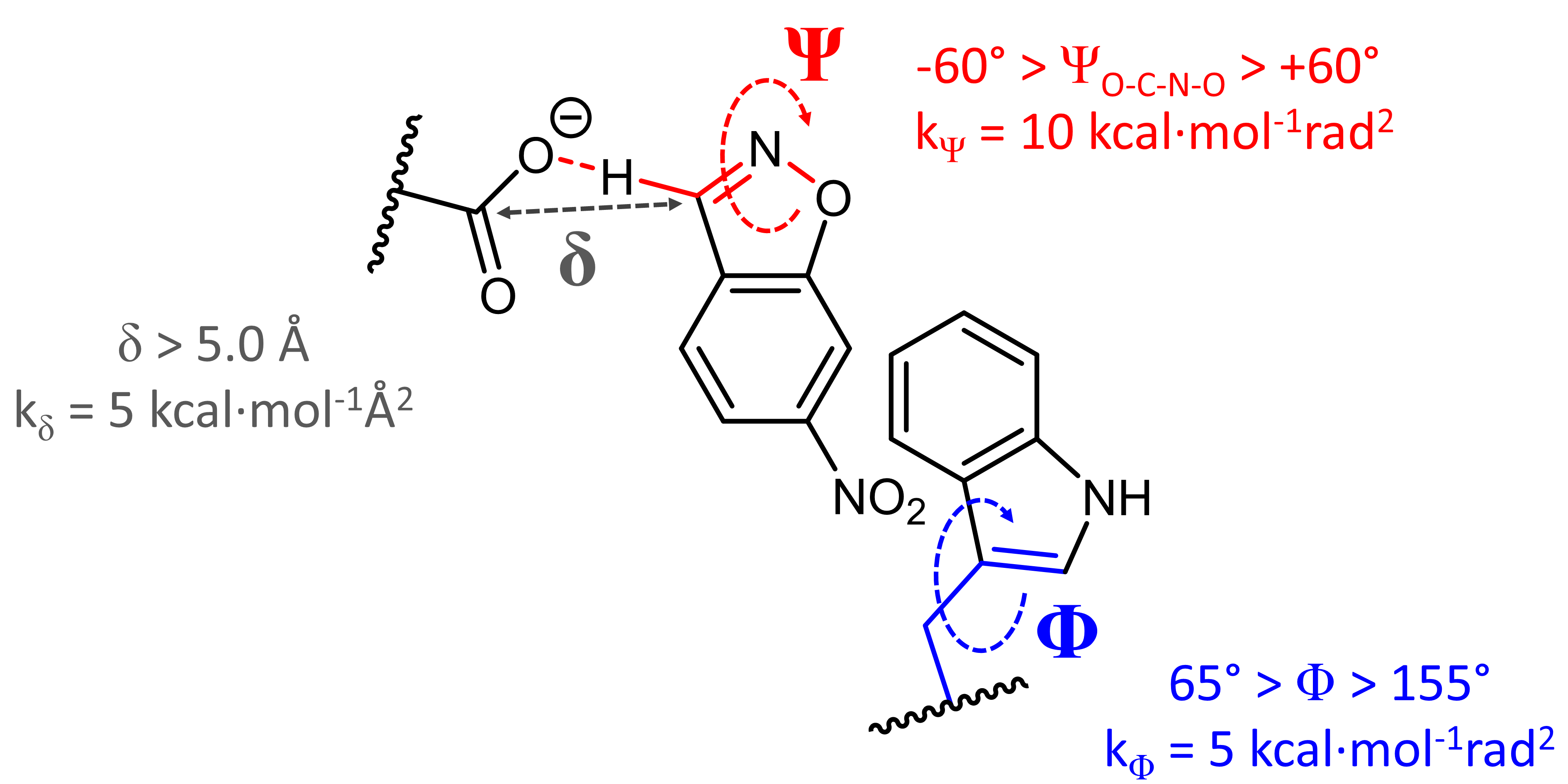

**Extended Data Fig. 4 | Restraints at the active site of the Kemp eliminases.** Similar to previous work,^1^ restraints were used to avoid moving away from conformations relevant for the reaction, owing to the long simulation timescale. Weak harmonic restraints for the ligand-base distance **δ** (black, >5.0 Å, 5.0 kcal∙mol^-1^Å^-1^) as well as the ligand-base dihedral angle **Ψ** (red, >±30°, 5.0 kcal∙mol^-1^rad^2^) were added to keep the ligand in the active site. Furthermore, the χ2 dihedral angle of Trp110 **Φ** (blue, >±60°, 5.0 kcal∙mol^-1^rad^-2^) was restrained, preventing its indole sidechain from blocking the active site. Importantly, the same restraints were added to all variants and ligands to avoid biasing the ${\Delta C}_{p}^{\ddagger}$ calculations.

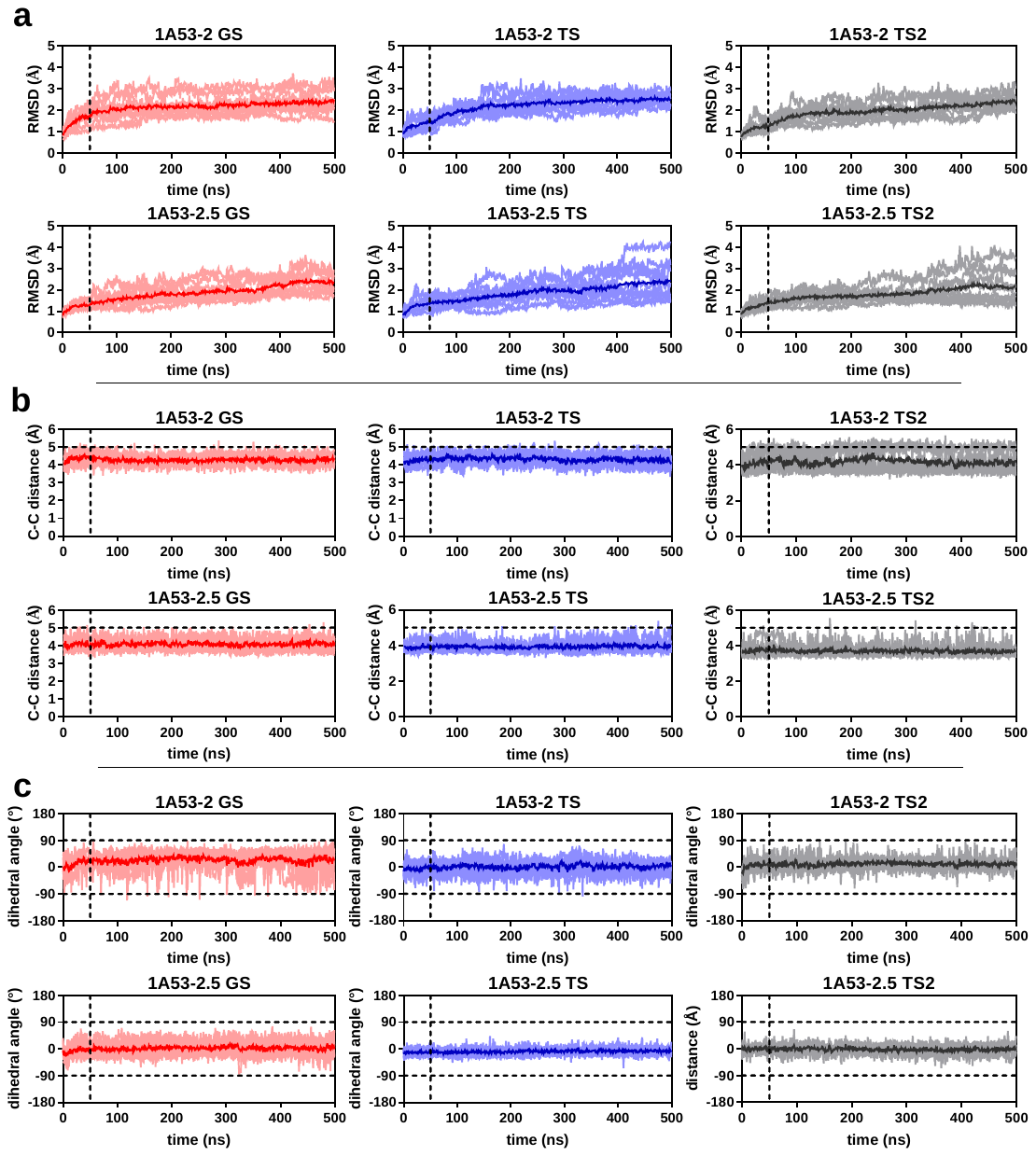

**Extended Data Fig. 5** **| MD trajectories.** Individual (light colours) and average (dark colours) trajectories of 1A53-2 and 1A53-2.5 with GS (red), TS (blue) and TS2 (grey) bound. **a,** Based on the RMSD plot, the first 50 ns of each simulations were discarded to avoid biasing of the analysis to the starting structures (dotted lines). The ligand-base distance **b,** and dihedral angle **c,** remained within the weak harmonic constraints (dotted lines) for most of the simulations. For a definition of the restraints see Extended Data Fig. 4.

|  | **without active-site waters** | |  | **with 10 active-site waters** | |
| --- | --- | --- | --- | --- | --- |
| **a** | 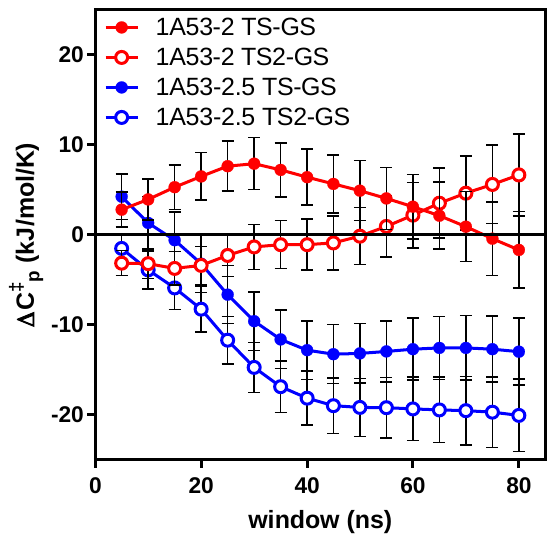 | |  | 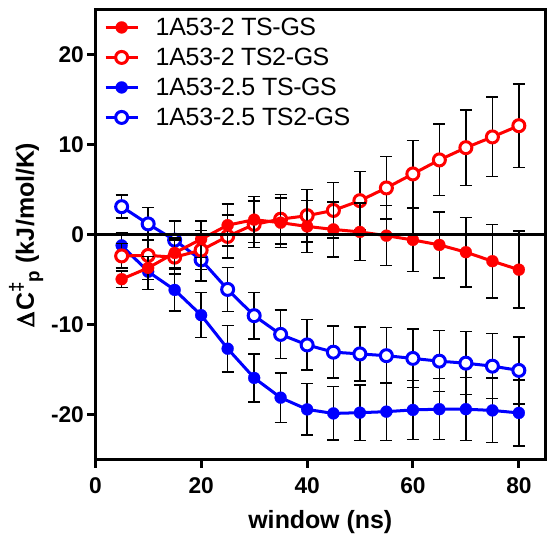 | |
| **b** | **1A53-2** | **1A53-2.5** |  | **1A53-2** | **1A53-2.5** |
| **GS** | 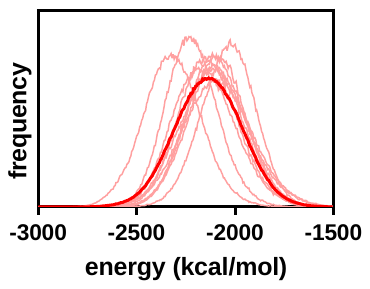 | 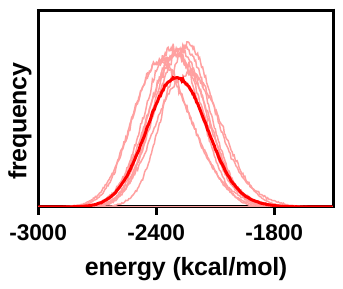 |  | 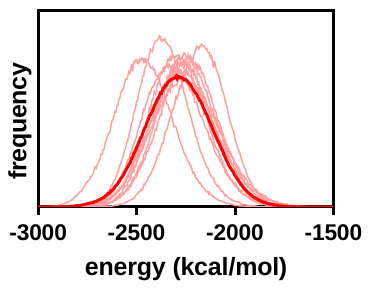 | 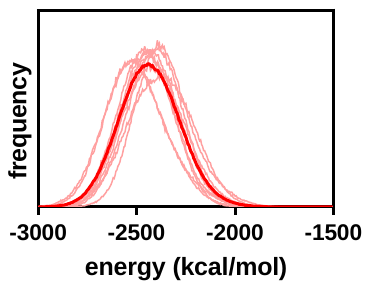 |
| **TS** | 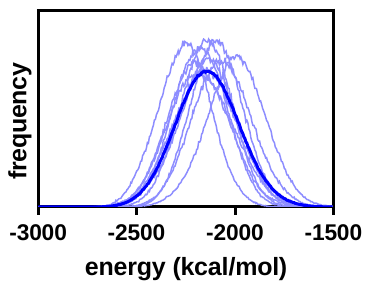 | 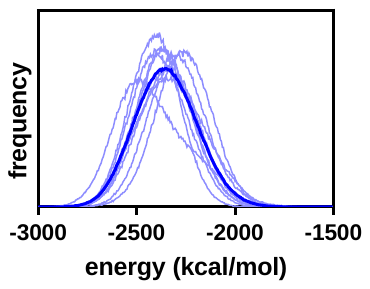 |  | 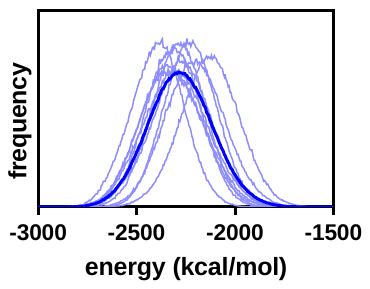 | 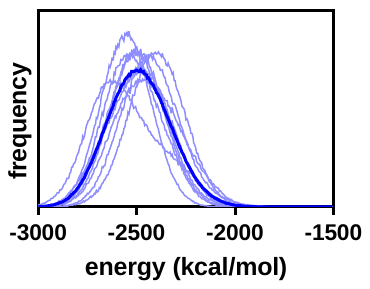 |
| **TS2** | 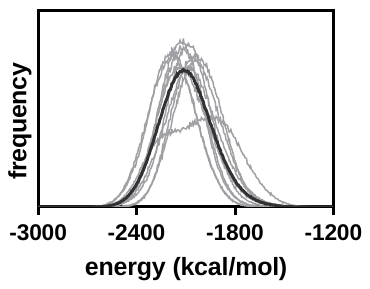 | 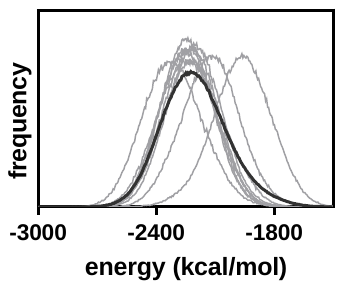 |  | 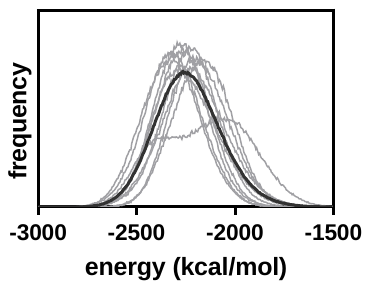 | 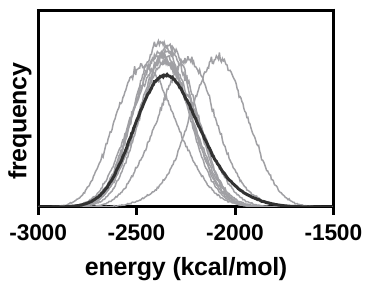 |

**Extended Data Fig. 6 |** **Energetic fluctuations give rise to** $\boldsymbol{\Delta C}_{\boldsymbol{p}}^{\boldsymbol{\ddagger}}$**.** Trajectories were analysed either dry (left), or with the 10 water molecules closest to Glu178 (right). **a,** Heat capacities were calculated from the difference in energetic variance of the transition and ground state for running windows with varying size (5 ns to 80 ns) for 1A53-2 (red) 1A53-2.5 (blue) for TS (filled circles) and TS2 (open circles). **b,** Energy distributions of the GS (red), TS (blue), and TS2 (grey) ensembles for ten individual trajectories (light colours) and their average (dark colours). The slow movement of some solvent exposed loops shifted the energy distributions in each trajectory. These slow movements become negligible for the short window sizes used for the analysis of ${\Delta C}_{p}^{\ddagger}$.

|  | **1A53-2** | | **0** | **1A53-2.5** | |
| --- | --- | --- | --- | --- | --- |
| **a** | **GS to TS** | | | | |
|  | 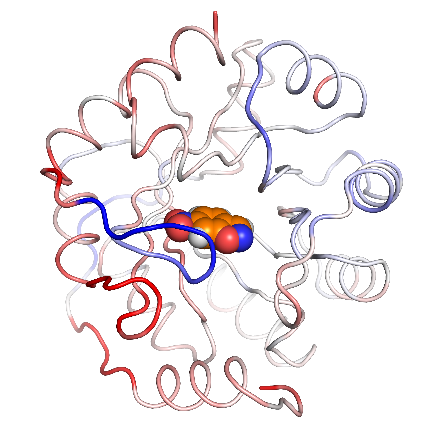 | 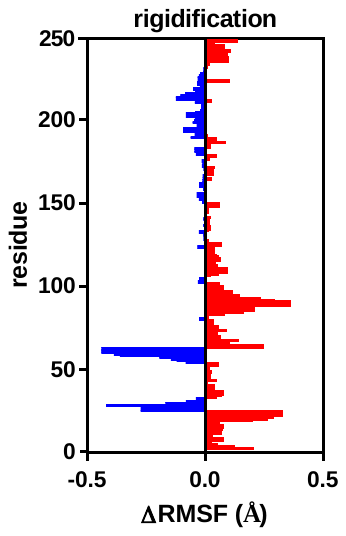 |  | 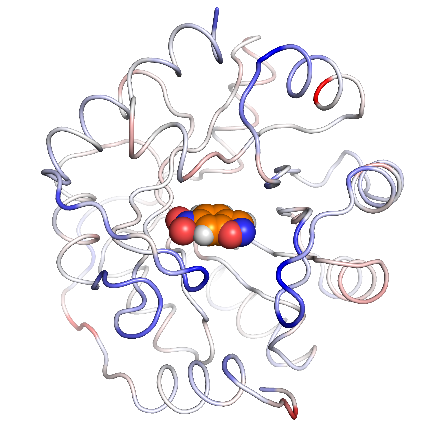 | 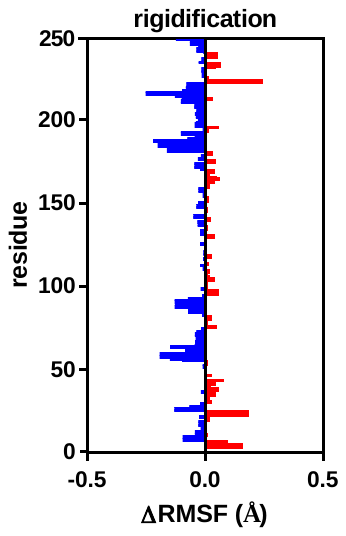 |
| **b** | **GS to TS2** | | | | |
|  | 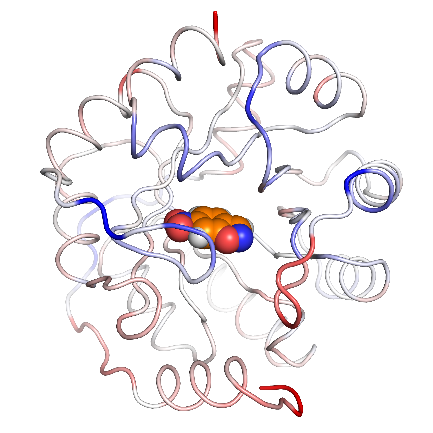 | 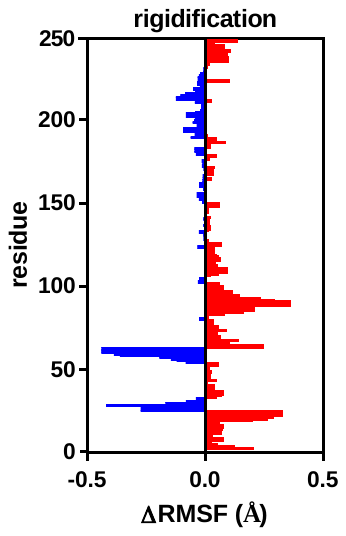 |  | 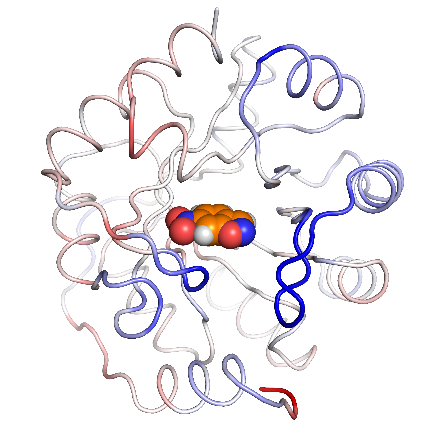 | 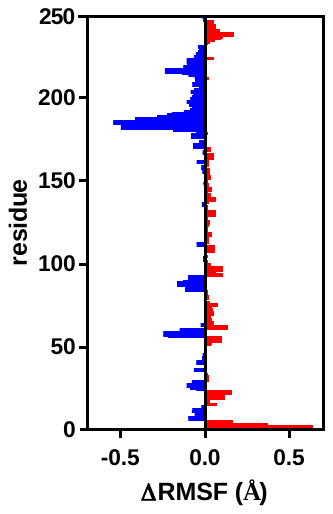 |

**Extended Data Fig. 7 | Structural fluctuations in 1A53-2 (left) and 1A53-2.5 (right).** Residues that rigidify (blue) or become more flexible (red) between **a,** GS and TS, and **b,** GS and TS2. In 1A53-2.5, rigidification is particularly expressed in three solvent exposed loops (residues 53-65, 84-92, and 181-192), which is reproduced by clustering based on the whole protein backbone. These loops were thus employed to cluster the structures in an open and closed state (Extended Data Fig. 8).

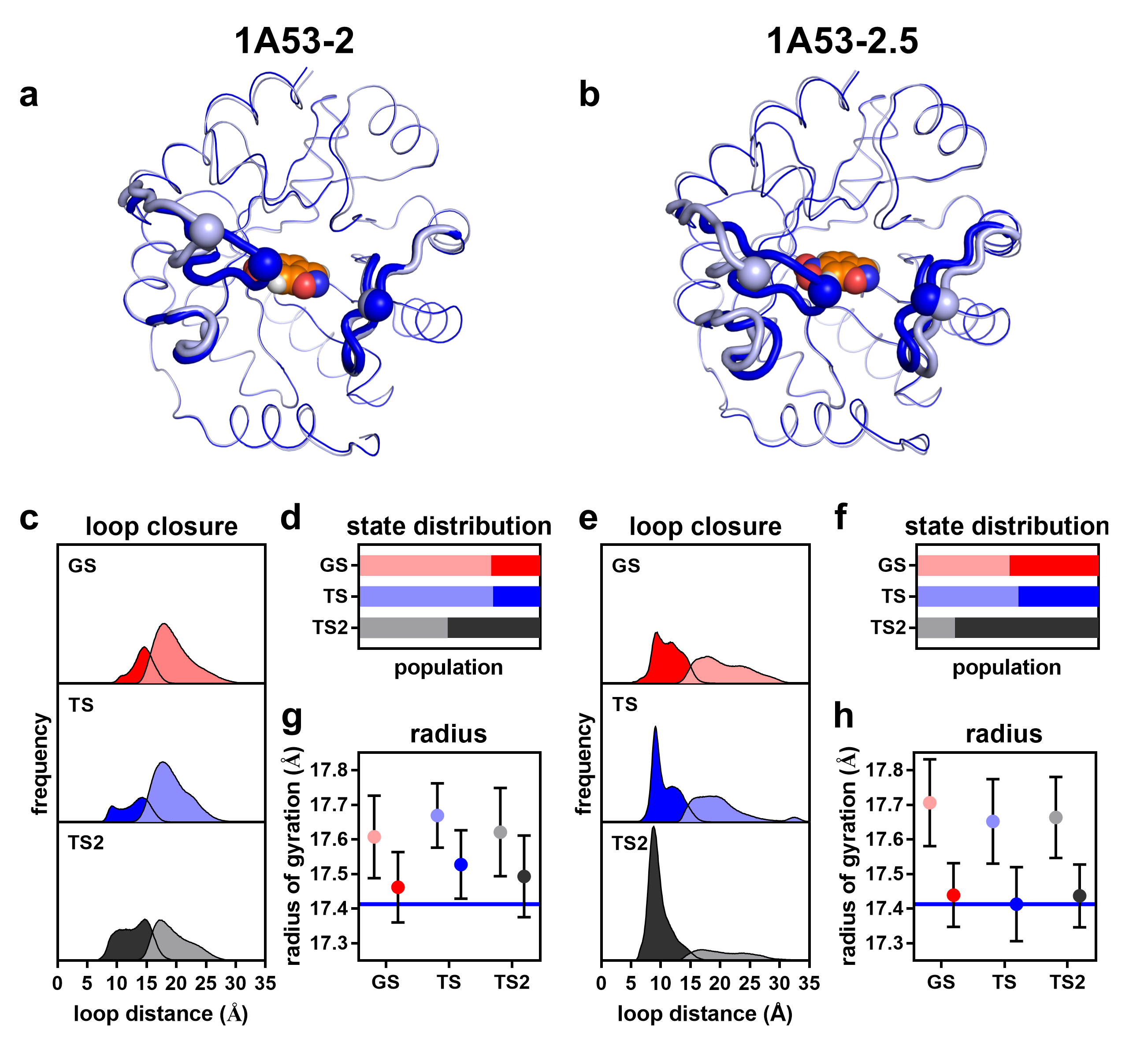

**Extended Data Fig. 8 | Cluster analysis.** Average structures of the **a,** 1A53-2 and **b,** 1A53-2.5 ensembles in the open (dark blue) and closed (light blue) state in complex with the TS. Distance-based clustering was performed using the k-means algorithm based on the three solvent exposed loops (bold tubes, residues 53-65, 84-92, and 181-192). Loop closure is indicated by the C_α_ distance of residues 58 and 188 (spheres). **c+e,** Histograms and **d+f,** distributions of the open (light colours) and closed (dark colours) states bound to GS (red), TS (blue) and TS2 (grey). **g+h,** Loop closure decreases the radius of gyration which became smallest in the closed TS of 1A53-2.5 (blue line).

|  | **1A53-2** | |  | **1A53-2.5** | |
| --- | --- | --- | --- | --- | --- |
| **a** | **GS** | | | | |
|  | 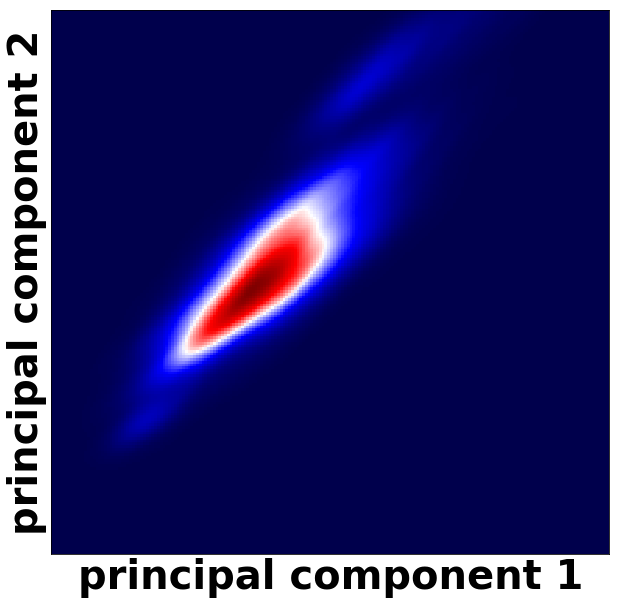 | 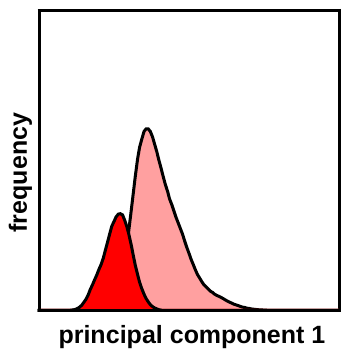 |  | 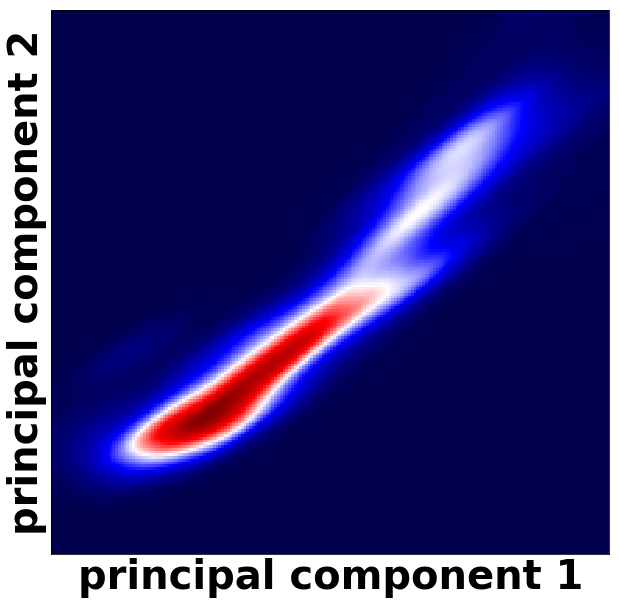 | 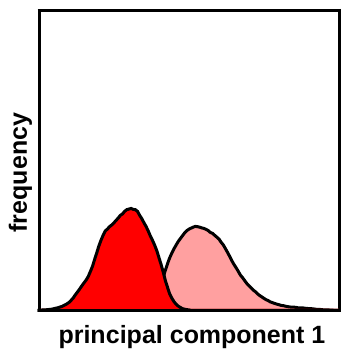 |
| **b** | **TS** | | | | |
|  | 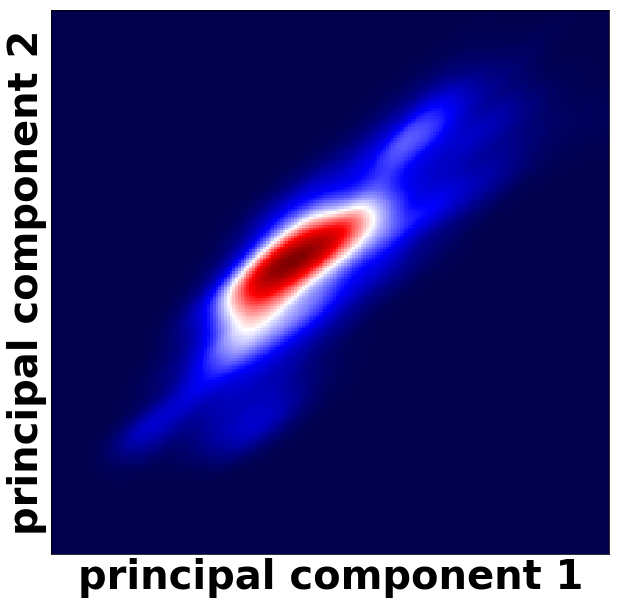 | 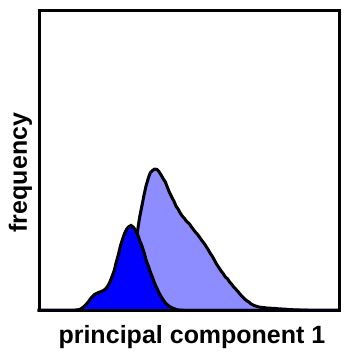 |  | 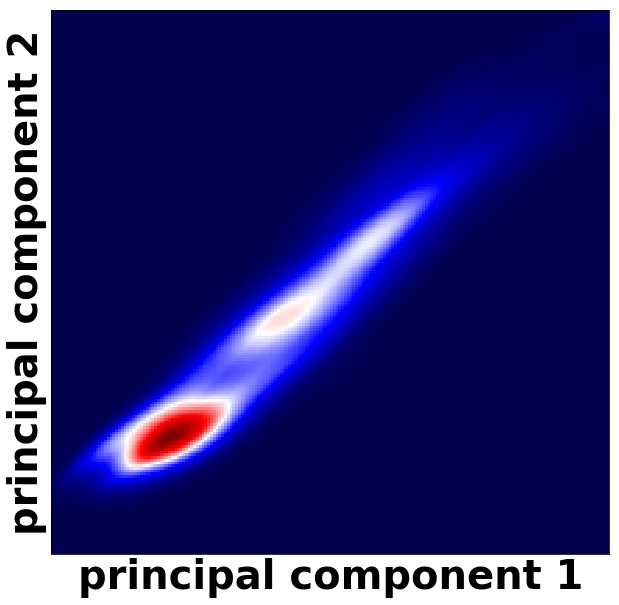 | 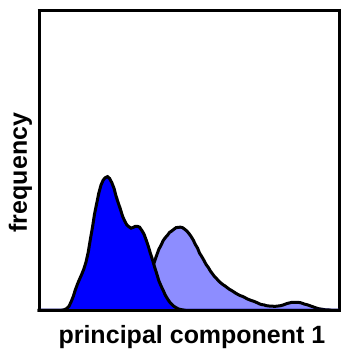 |
| **c** | **TS2** | | | | |
|  | 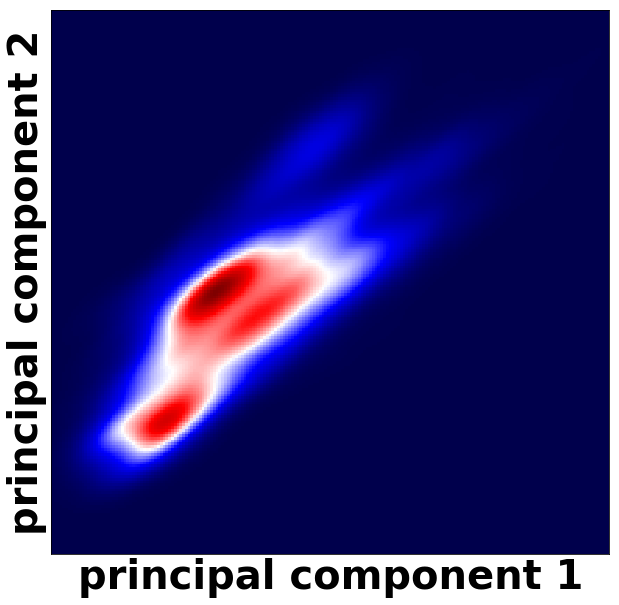 | 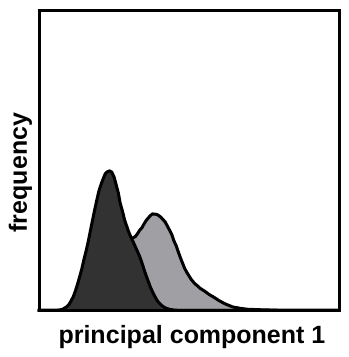 |  | 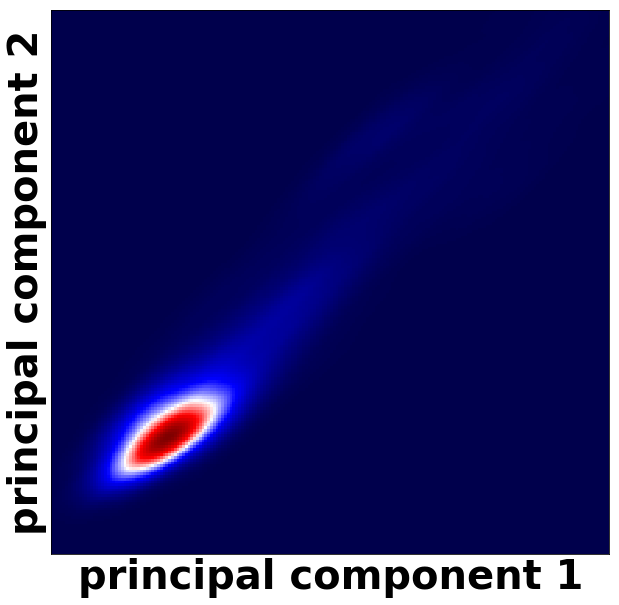 | 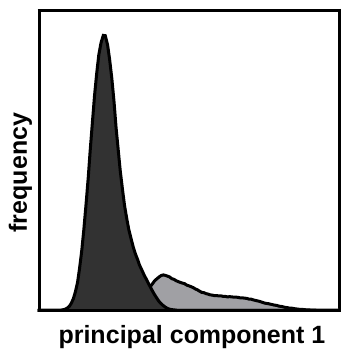 |

**Extended Data Fig. 9 |** **Principal component analysis reproduces two state model.** Principal component analysis of the mobile loops (residues 53-65, 84-92, and 181-192) was performed for the **a,** GS (red), **b,** TS (blue), and **c,** TS2 (grey) ensembles of 1A53-2 (left) and 1A53-2.5 (right). The histogram of the first principal component after partition of the trajectory according to the cluster analysis (Extended Data Fig. 8) into open (light) and closed (dark) states reproduces the two state model.

|  | **1A53-2** | |  | **1A53-2.5** | |
| --- | --- | --- | --- | --- | --- |
|  | **GS to TS** | **GS to TS2** |  | **GS to TS** | **GS to TS2** |
| **a** | 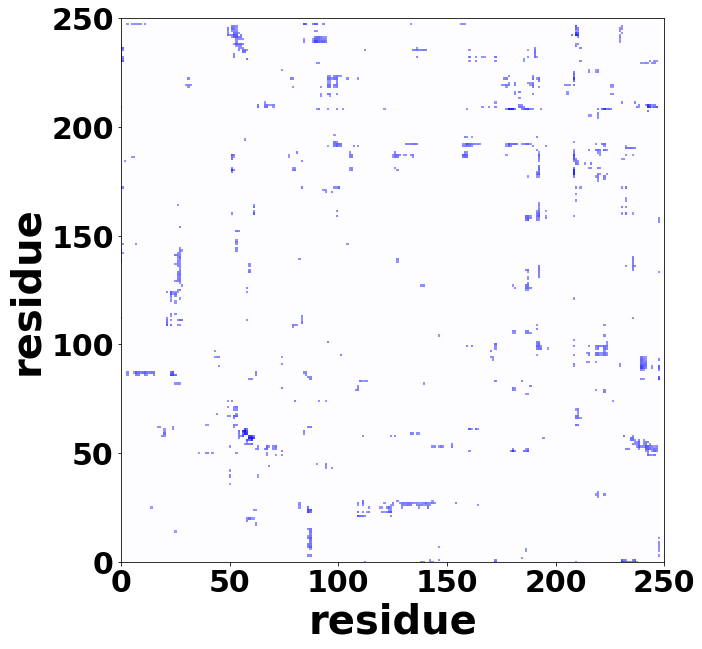 | 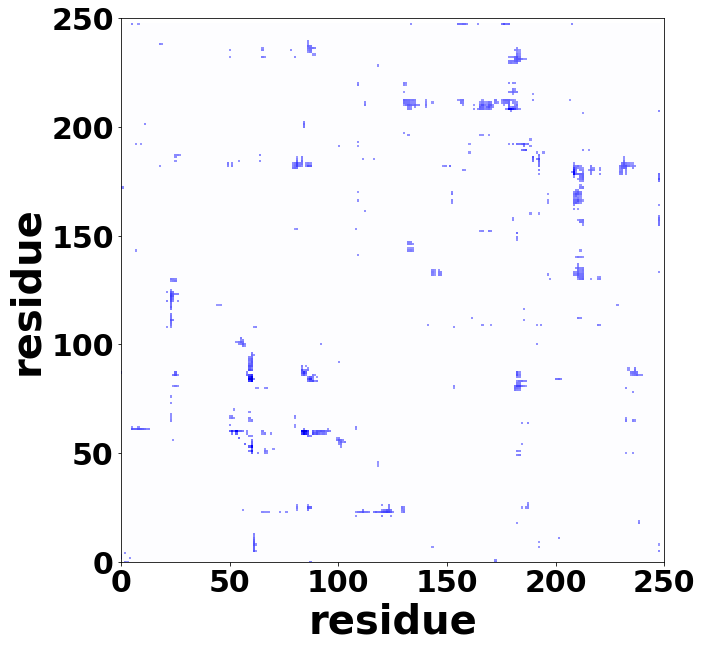 |  | 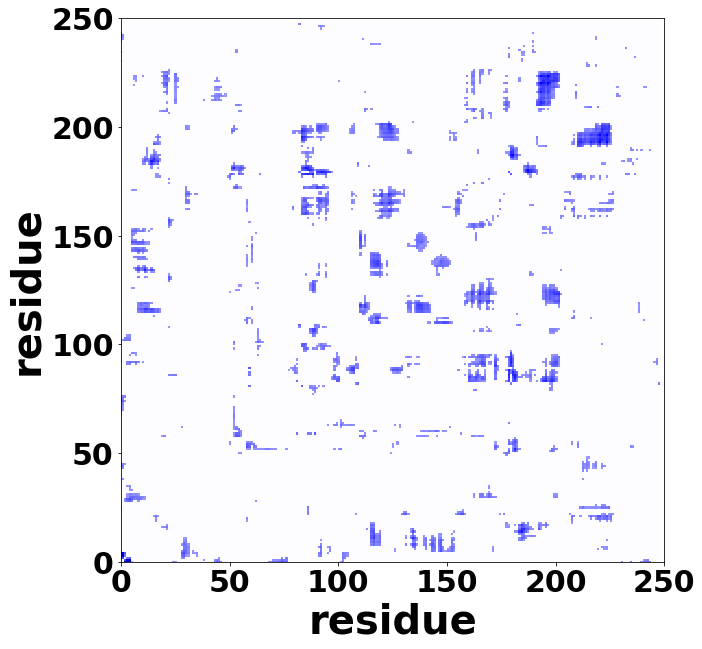 | 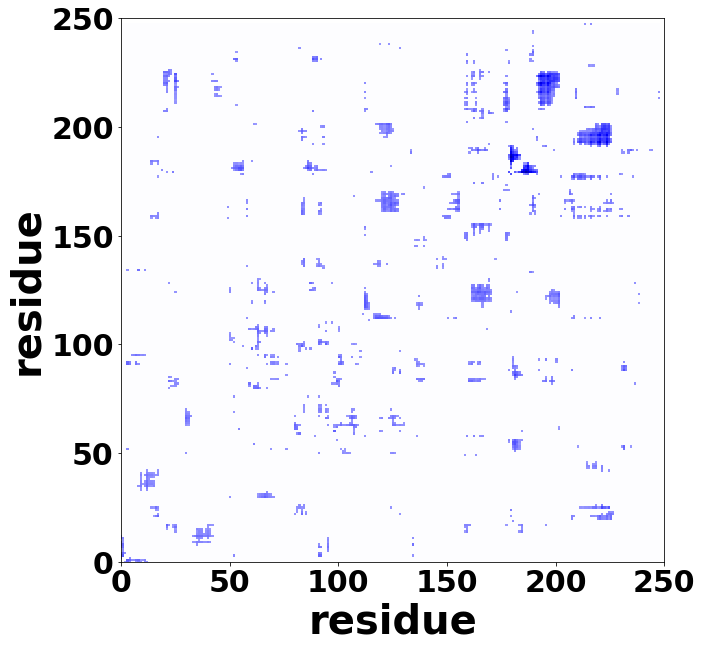 |
| **b** | 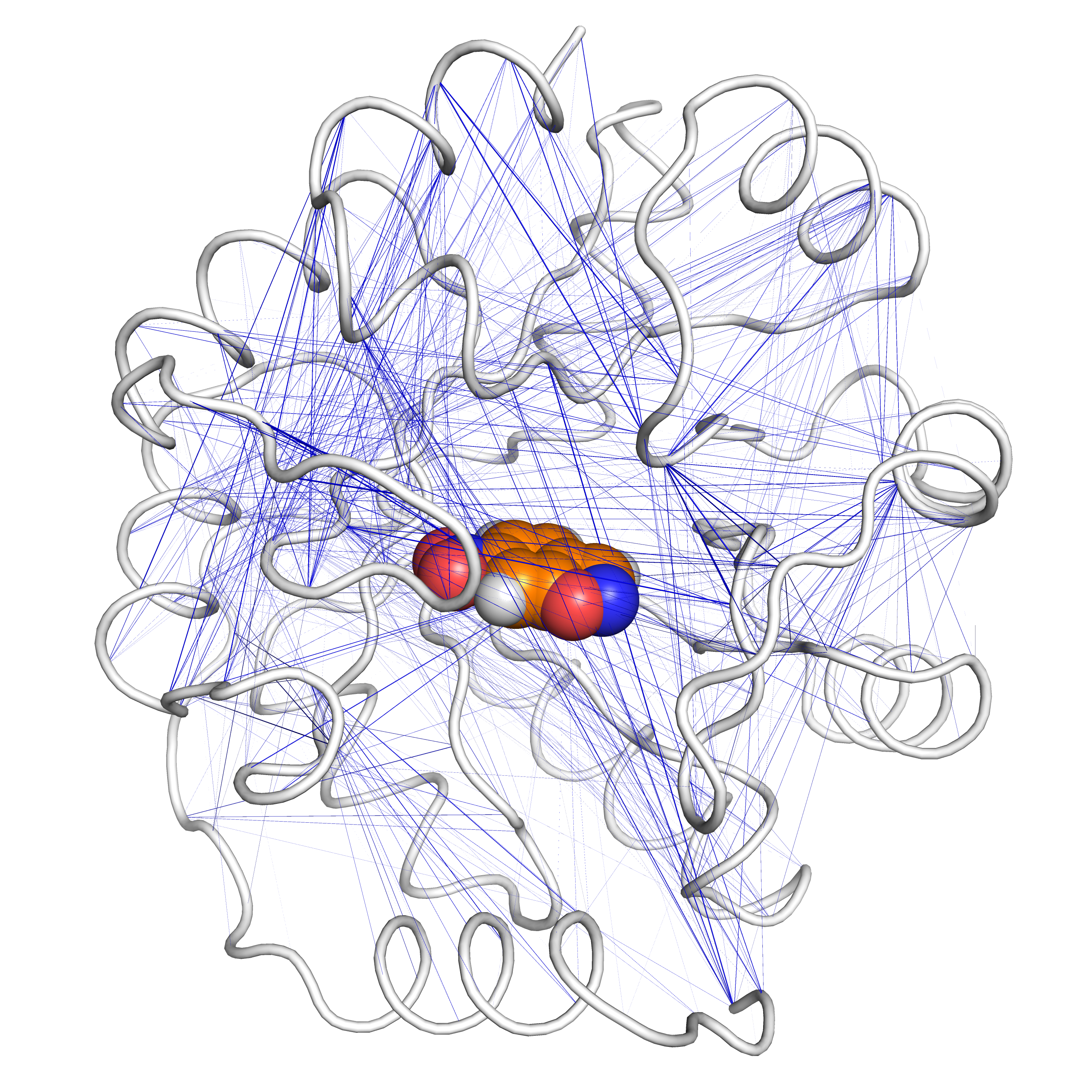 | 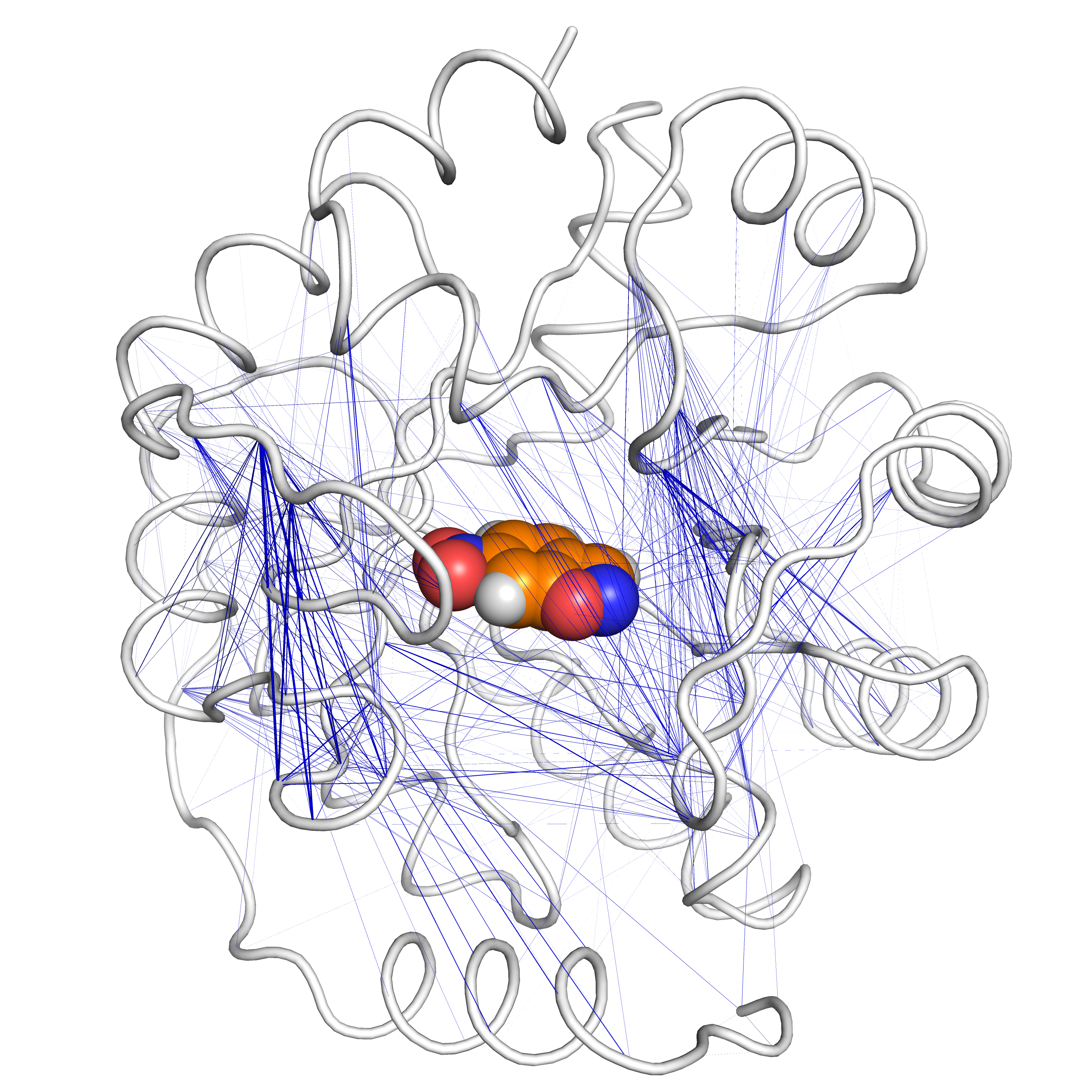 |  | 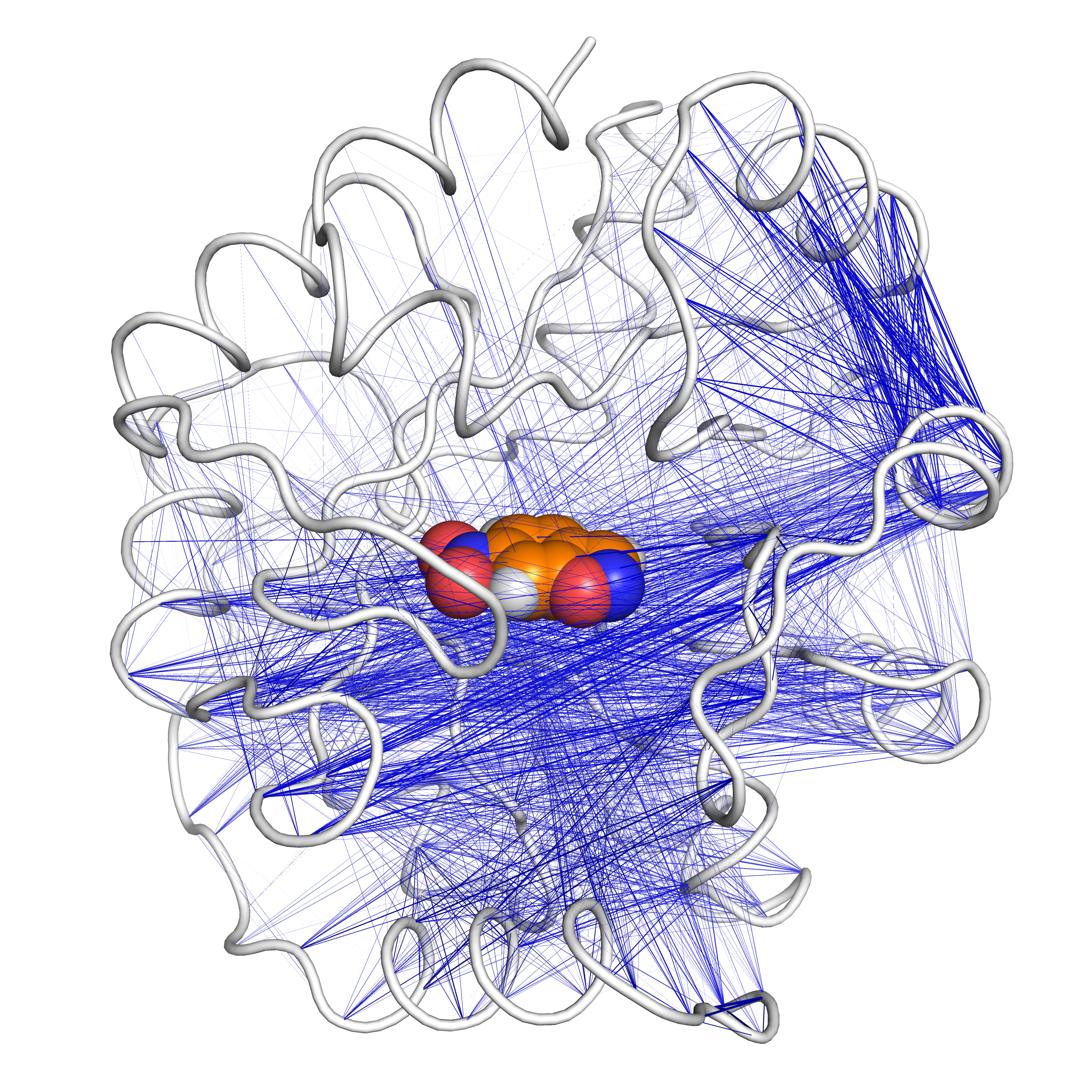 | 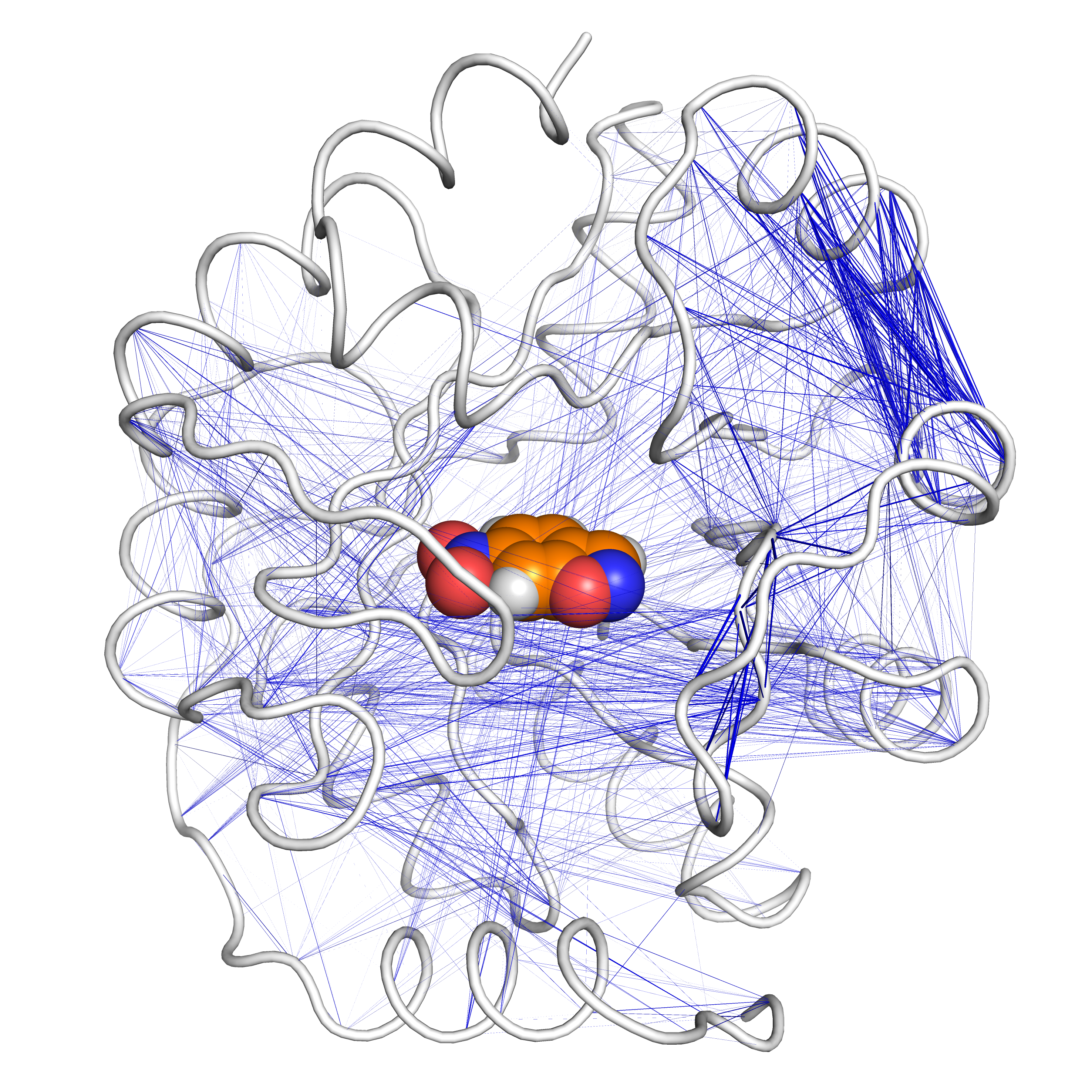 |
| **c** | 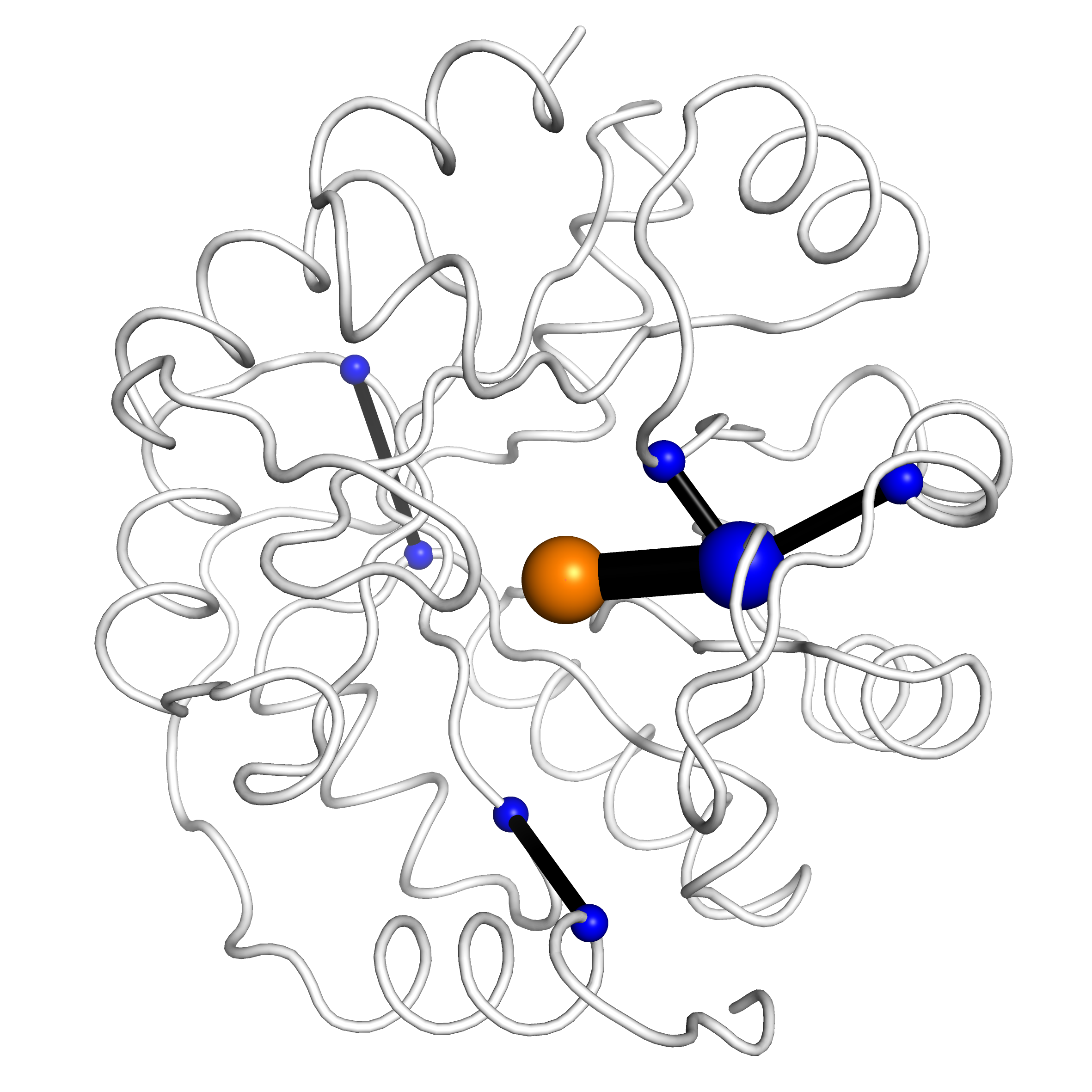 | 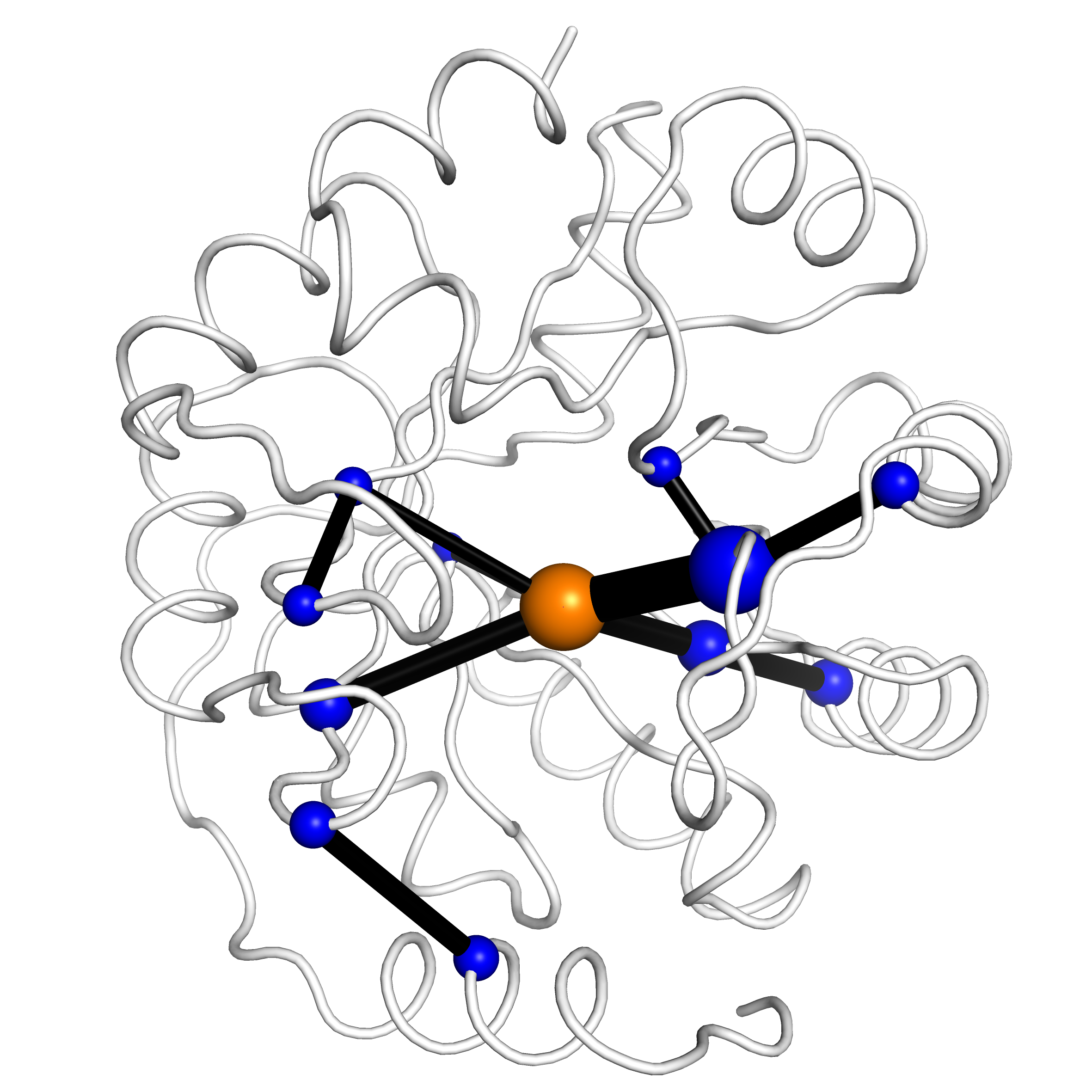 |  | 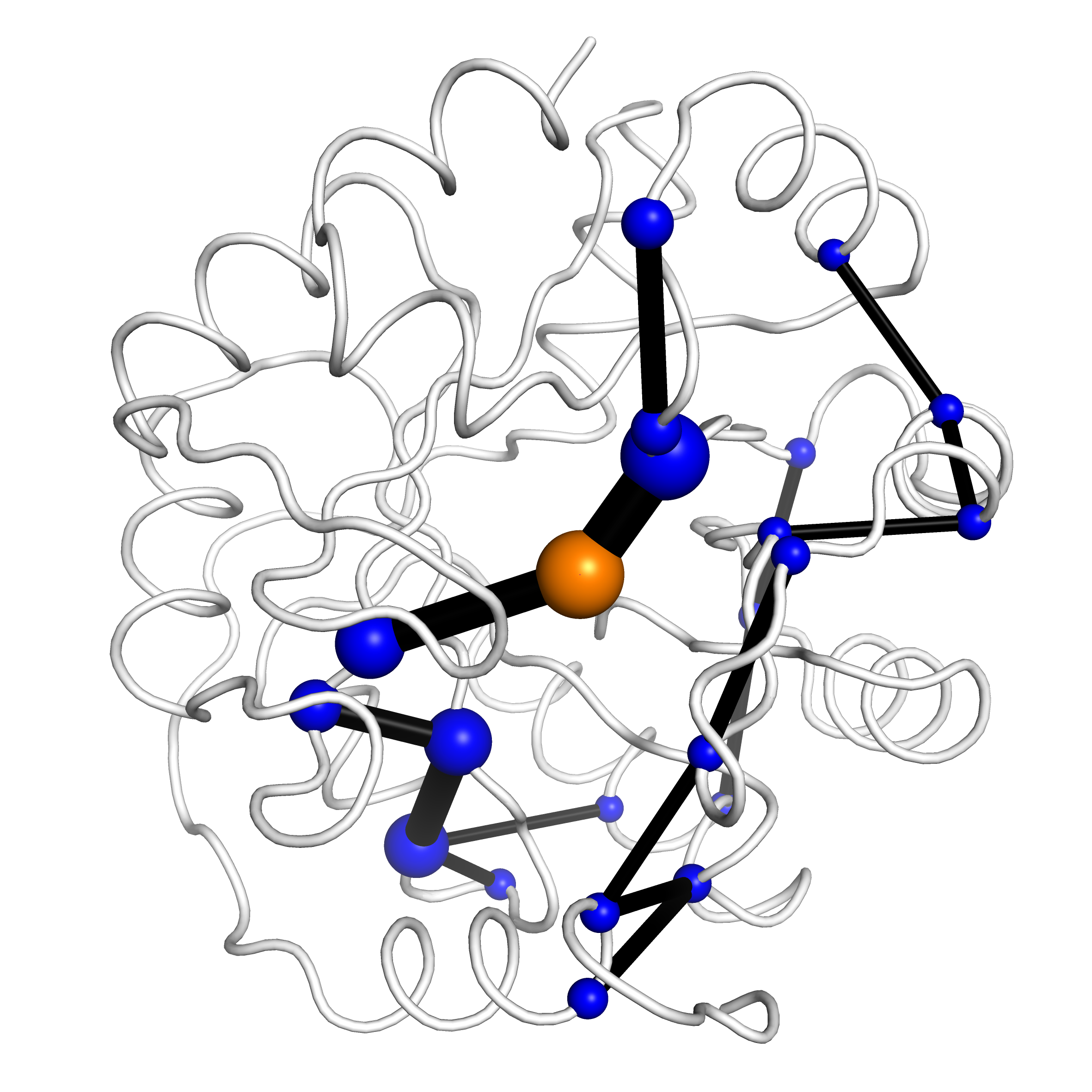 | 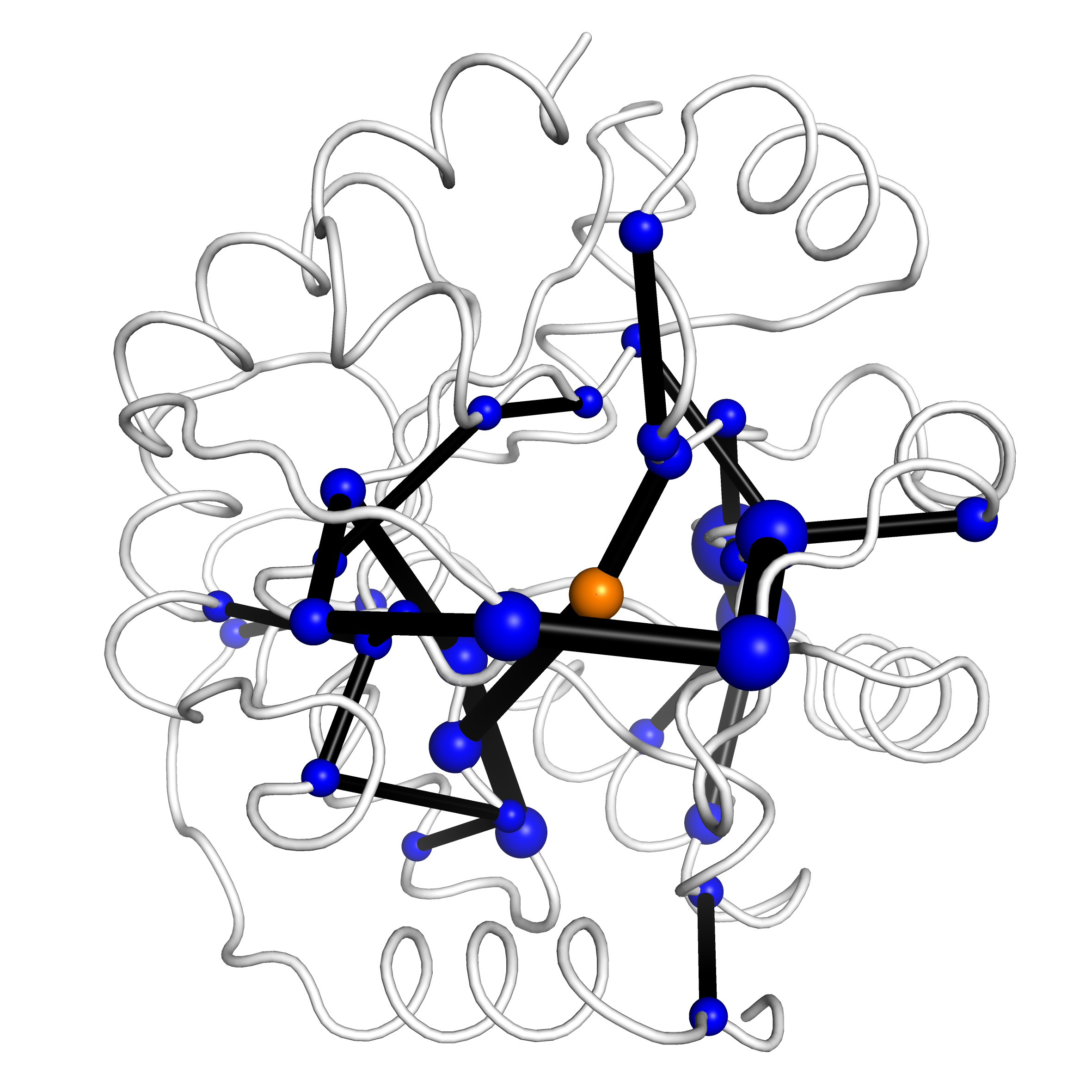 |

**Extended Data Fig. 10** **| Dynamical networks arising in the closed ensembles of 1A53-2 (left) and 1A53-2.5 (right) between GS and either TS or TS2.** **a**, Backbone movements become highly correlated in the closed TS ensemble of 1A53-2.5, indicating the presence of global vibrations comprising most of the protein. **b,** Cross correlations that increase between GS and TS by ≥20% are indicated as lines on the structures. **c,** Based on the cross-correlations, shortest pathway maps were calculated^19^. These show that evolution introduced a communication network in 1A53-2.5 that originates from the TS and comprises vast parts of the closed state. The size of the edge (black sticks) and vertices (blue spheres: protein, orange sphere: ligand) indicate the increasing weight of the network between GS and TS.

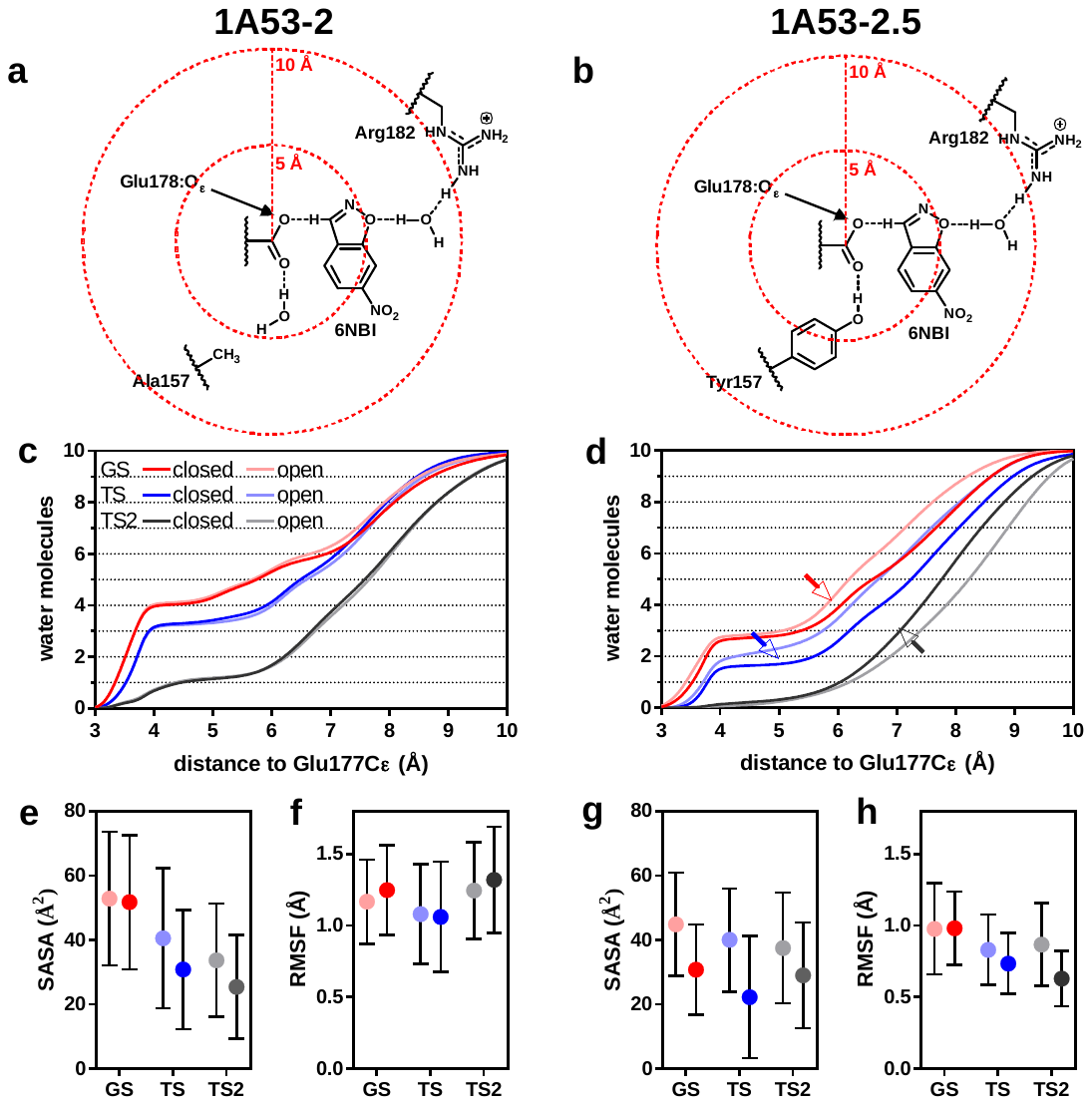

**Extended Data Fig. 11** **| Local effects at the active sites of** **1A53-2 (left) and 1A53-2.5 (right).** **c+d,** The distance-distribution of the ten water molecules closest to the O_ε_ of Glu178 is shown for the GS (red), TS (blue), and TS2 (grey) in the closed (dark colours) and open (light colours) state. The active site becomes less hydrated in the transition states compared to the ground state for both enzymes. Furthermore, the A157Y mutation in 1A53-2.5 displaces a water molecule in the evolved variant at approximately 3 Å. Remarkably, loop closure expels water from the active site in 1A53-2.5 (arrows) which was not observed for 1A53-2. The **e+g,** SASA of the ligand and **f+h,** RMSF of the ligand and base are furthermore lower in 1A53-2.5 than 1A53-2, indicating tighter packing of the ligand.
