## Supplementary material for "Evolution of dynamical networks enhances catalysis in a designer enzyme": Methods

Methods for

**Model construction.** Ground state (Michaelis complex) and transition state geometries were determined with the GAUSSIAN 16 quantum chemistry software package^1^. The models comprised 6-nitrobenzisoxazole and acetate and were optimized at the B3LYP/6-311++g(d,p) level; other methods gave similar results. The transition state was located as a maximum on the potential energy surface for the cleavage of the scissile C-H bond by performing a relaxed distance scan of this bond, similar to previous work^2^. Transition state geometries were subsequently optimized using the Berny algorithm. CHELPG charges (MP2/6-311g++(d,p)) were determined for each state (Extended Data Fig. 1). Ground state (GS) and transition state (TS) models were constructed based on these charges and the General Amber Force-Field (GAFF). An additional transition state model (TS2) was constructed with the same geometry and charge distribution as TS, but with the proton bound to Glu178 instead of to the ligand. Input structures for MD were manually constructed by replacing the 6-nitrobenzotriazole inhibitor in the crystal structures of 1A53-2 (PDB code: 3NZ1)^3^ and 1A53-2.5 (PDB: 6NW4)^4^ with either the substrate or transition state model. Different forcefields were tested for representing the proteins. For MD simulations with the CHARMM36 **protein** force-field, the CHARM-GUI was used to parametrize the protein^5^, and to solvate and neutralize the complex in an octahedral box with a 10 Å edge distance in TIP3P water and 0.15 M sodium chloride. For simulations with the **AMBERff99SB protein force-field,** tLEAP was used to solvate the complex in an octahedral box with a 10 Å edge distance in TIP3P water and to neutralize with chloride ions^6^. All MD simulations were run with the Amber16 package^6^.

**Testing force-fields with MM and QM/MM. The CHARMM36 and AMBERff99SB protein force-fields were tested in MM and QM/MM MD simulations. Simulations with the CHARMM36 force-field gave structures and relative activation energies consistent with experiment. While the GS complex of 1A53-2 was not stable with AMBERff99SB, which apparently underestimated the π-stacking interactions of the GS with Trp110 and Trp210 leading to displacing of the ligand with water within <100 ns, CHARMM36 gave structures consistent with the experimentally observed structures, and allowed sampling on the timescales required for the analysis. The finding that CHARMM36 more accurately describes the bound ligands is mimicked by hybrid QM/MM simulations, in which CHARMM36 reproduced the activity improvements during evolution. To that end, the distances of the scissile C-H and N-O bonds were sampled** similar to previously published methods^7,8^. **The QM subsystem was composed of the substrate and the carboxylate and γ-methylene of Glu178 and was described using either the semiempirical AM1 or PM6 Hamiltonian. To saturate the valence of the QM/MM frontier atoms, a link atom was placed between the C_β_ and C_γ_ atoms of Glu178. A 15** Å **cut-off for nonbonding interactions with the QM subsystem was applied. QM/MM MD was started from structures that were equilibrated with the TS model (see MD simulations). After 1 ns equilibration with 1 fs time steps, umbrella sampling of the scissile C-H and N-O bonds was started. C-H and N-O bonds were sampled in 0.1** Å **and 0.05** Å **steps with force constants of 200 kcal∙mol**^-1^Å^-1^ **and 400 kcal∙mol**^-1^Å^-1^, respectively**. Each window was equilibrated by ramping up the force constants from 0 to their final value over 10 ps and simulation over additional 10 ps. Subsequently, 40 ps of sampling were performed. Potential of mean force free-energy landscapes were calculated from these data using the weighted histogram analysis method^9^. The P**M6/CHARMM36 and **AM1**/CHARMM36 QM/MM simulations correctly reproduced the activation energy difference between **1A53-2 than 1A53-2.5**, while PM6/**AMBERff99SB gave higher activation energies for 1A53-2 than 1A53-2.5 (**Extended Data Fig. 3), probably because interactions of the ligand with the active site (e.g. **π-stacking to Trp110 and Trp210) were not accurately reflected in AMBERff99SB**. Thus, CHARMM36 was employed for all subsequent simulations.

**MD simulations.** All simulations were performed using established and validated protocols^10^. Briefly, the structures were minimized by 300 step minimization of water, ions and hydrogens only, followed by 50 ps MD simulation in the NVT ensemble at 300 K of the water and ions only (positional restraint on protein atoms and ligand: 25 kcal·mol^‑1^·Å^-2^), and minimization of the whole system with positional restraints on C_α_ atoms (100 kcal·mol^-1^·Å^-2^) for 300 steps. Ten independent simulations were performed for each protein and ligand combination by assigning random velocities at 25 K followed by heating to 300 K in 20 ps using Langevin dynamics for temperature control with a 1 ps^−1^ collision frequency (maintaining 100 kcal·mol^-1^·Å^-2^ positional restraints on C_α_ atoms). For each independent simulation, four consecutive 10 ps simulations were subsequently performed under the same conditions, reducing the positional restraints on the C_α_ atoms to 20, 8, 4, 3, 2, and 1 kcal·mol^-1^·Å^-2^. Subsequently, 1 ns of equilibration was performed in the NPT ensemble at 1 atm, using the Berendsen barostat (10 ps pressure relaxation time) and Langevin dynamics for temperature control (1 ps^−1^ collision frequency). Production simulations (500 ns) were performed in the NPT ensemble with the Berendsen thermostat and loose temperature coupling and pressure scaling (10 ps time constant each), to limit the influence of the thermostat while avoiding temperature drift. In all MD simulations, the default direct-space cutoff was used for non-bonded interactions, with particle-mesh Ewald summation for long-range electrostatic interactions. MD simulations were run with pmemd.cuda on GPUs, using the SPFP precision model^12^. Throughout the simulations, restraints were used to maintain the Michaelis complex, with equivalent restraints in each state (see Extended Data Fig. 4 for a detailed description), similar to previous simulations^10^.

**Heat capacity calculations.** Analysis of ${\Delta C}_{p}^{\ddagger}$ was performed using procedures developed and tested in previous work on natural enzymes^10^, based on structures sampled at 10 ps intervals from 50–500 ns windows of the MD simulations, with force-field energies recalculated after stripping of ions and all but 10 water molecules closest to the catalytic base. Energy distributions were calculated by normalization to a moving average (50 ns window). Heat capacities were similarly calculated based on the moving average of the energy variance according to equation 1, where ${\Delta C}_{A-B}$ is the change in heat capacity between two states, A and B with the variances $\left\langle{\delta H}_{A}^{2} \right\rangle$ and $\left\langle{\delta H}_{B}^{2} \right\rangle$. Error-bars were obtained by leave-one-out cross-validation.

${\Delta C}_{A-B}=\frac{\left\langle{\delta H}_{A}^{2} \right\rangle-\left\langle{\delta H}_{B}^{2} \right\rangle}{k_{B}T^{2}}$	(1)

**Clustering and principal component analysis.** All other analyses were performed with CPPTRAJ^6^, unless otherwise stated. RMSF calculations were performed for the backbone C_α_ atoms of each trajectory, and RMSF values represent the average of the ten independent trajectories. Clustering was performed using the k-means algorithm. Initial clustering based on the whole protein backbone revealed that structural changes are predominantly located in three solvent exposed loops (residues 53-65, 84-92, and 181-192), in agreement with the RMSF calculations. The final cluster analysis presented here was performed for these loops in distance-space with the k-means algorithm. Similarly, the distance-matrix of these loops was analysed by principal component analysis using MDtraj^11^, and the first principal component aligned well with the two states from the cluster model. For the following analysis, each MD trajectory was partitioned into open and closed state based on the cluster model.

**Calculation of local effects.** Interaction energies were calculated for each protein residue with the ligand using Sander^6^. SASA and RMSF were calculated for the ligand and base only, after alignment of the C_α_ in the whole trajectory. All values were individually calculated for the ten independent trajectories and subsequently averaged.

**Dynamic cross correlations and shortest pathway maps.** Networks were calculated using established protocols^12^. Briefly, dynamic cross correlations and distance matrices were calculated with CPPTRAJ based on the backbone C_α_ positions and the ligand centre of mass^6^. Dynamical networks were calculated based on the difference in cross correlations between the transition state and ground state. The distance cut-off was increased to 10 Å, beyond that used in previous work^12^, to ensure that all active site residues can interact with the ligand.

**References**

1	Frisch, M. J. *et al.* Gaussian 16. *Wallingford, CT* (2016).

2	Na, J., Houk, K. N. & Hilvert, D. Transition state of the base-promoted ring-opening of isoxazoles. Theoretical prediction of catalytic functionalities and design of haptens for antibody production. *J. Am. Chem. Soc.* **118**, 6462-6471 (1996).

3	Privett, H. K. *et al.* Iterative approach to computational enzyme design. *Proc. Natl. Acad. Sci. U. S. A.* **109**, 3790-3795 (2012).

4	Bunzel, H. A. *et al.* Emergence of a negative activation heat capacity during evolution of a computationally designed enzyme. *J. Am. Chem. Soc.* **141**, 11745-11748 (2019).

5	Jo, S., Kim, T., Iyer, V. G. & Im, W. CHARMM-GUI: A Web-based Graphical User Interface for CHARMM. *J. Comput. Chem.* **29**, 1859-1865 (2008).

6	Case, D. A. *et al.* AMBER 2016. *University of California, San Francisco* (2016).

7	Alexandrova, A. N., Röthlisberger, D., Baker, D. & Jorgensen, W. L. Catalytic Mechanism and Performance of Computationally Designed Enzymes for Kemp Elimination. *J. Am. Chem. Soc.* **130**, 15907-15915 (2008).

8	Swiderek, K., Tunon, I., Moliner, V. & Bertran, J. Revealing the origin of the efficiency of the de novo designed Kemp eliminase HG-3.17 by comparison with the former developed HG-3. *Chemistry* **23**, 7582-7589 (2017).

9	Grossfield, A. WHAM: the weighted histogram analysis method, version 2.0.9. *http://membrane.urmc.rochester.edu/wordpress/?page_id=126* (2019).

10	van der Kamp, M. W. *et al.* Dynamical origins of heat capacity changes in enzyme-catalysed reactions. *Nat. Commun.* **9**, 1177 (2018).

11	McGibbon, Robert T. *et al.* MDTraj: A Modern Open Library for the Analysis of Molecular Dynamics Trajectories. *Biophys. J.* **109**, 1528-1532 (2015).

12	Osuna, S., Jimenez-Oses, G., Noey, E. L. & Houk, K. N. Molecular dynamics explorations of active site structure in designed and evolved enzymes. *Acc. Chem. Res.* **48**, 1080-1089 (2015).
